# Induction of antiviral Interferon-Stimulated Genes (ISGs) by neuronal STING promotes the resolution of pain

**DOI:** 10.1101/2023.08.04.551993

**Authors:** Manon Defaye, Amyaouch Bradaia, Nasser S. Abdullah, Francina Agosti, Mircea Iftinca, Vanessa Soubeyre, Kristofer Svendsen, Gurveer Gill, Mélissa Cuménal, Nadine Gheziel, Jérémy Martin, Gaetan Poulen, Nicolas Lonjon, Florence Vachiery-Lahaye, Luc Bauchet, Lilian Basso, Emmanuel Bourinet, Isaac M. Chiu, Christophe Altier

## Abstract

Inflammation and pain are intertwined responses to injury, infection, or chronic diseases. While acute inflammation is essential in determining pain resolution and opioid analgesia, maladaptive processes occurring during resolution can lead to the transition to chronic pain. Here we found that inflammation activates the cytosolic DNA-sensing protein Stimulator of Interferon Genes (STING) in DRG nociceptors. Neuronal activation of STING promotes signaling through TANK-binding kinase 1 (TBK1) and triggers an interferon-beta (IFNβ) response that mediates pain resolution. Notably, we found that mice expressing a nociceptor-specific gain-of-function mutation in STING exhibited an IFN gene signature that reduced nociceptor excitability and inflammatory hyperalgesia through a KChIP1-Kv4.3 regulation. Our findings reveal a role of IFN-regulated genes (IRGs) and KChIP1 downstream of STING, in the resolution of inflammatory pain.

## Introduction

Pain and inflammation are two integrated biological responses that serve as the main defense mechanisms during injury or infection. Inflammatory pain is adaptive and protective, however it often persists even after the inflammation has subsided. While it is well established that activated immune cells contribute to acute pain ^1,2^, recent studies suggest that acute inflammation can also be protective against the transition to chronic pain ^3^. Therefore, understanding how pain resolves during inflammation is crucial for preventing the development of chronic pain.

Like the immune system, the nociceptive system alerts the host to the presence of “danger signals” ^4^, and certain pathogens have evolved mechanisms to disable sensory modalities, such as smell, taste ^5–7^ and even pain ^6^, by targeting danger signal receptors. Nociceptors have the ability to respond to pathogen/damage-associated molecular patterns (PAMPs/DAMPs) through various pattern recognition receptors (PRRs) ^8^. Toll-like receptors (TLRs), Nod-like receptors (NLRs), RIG-I-like receptors (RLRs), and cytosolic DNA sensors (CDSs) have been reported to be expressed in dorsal root ganglion neurons ^8^.

Among these receptors, Stimulator of Interferon Genes (STING) is a cytosolic DNA sensor that plays a role in recognizing self-DNA, viral DNA, and cyclic dinucleotides (CDN: c-di-GMP, 3’3’-cGAMP and c-di-AMP) produced by bacteria. STING activation leads to the production of Type I interferons (IFN-I) such as interferon α (IFN α) and interferon β (IFNβ) isoforms. Bacterial and host-derived CDNs induce STING dimerization, which activates TANK Binding Kinase 1 (TBK1). TBK1 phosphorylates interferon regulatory factor 3 (IRF3), leading to the production of IFN-I and other genes involved in host responses, promoting elimination of pathogens and damaged host cells upon inflammation ^9^. IFN-I bind to IFNα/β receptor (IFNAR) on producer and nearby cells. This autocrine and paracrine stimulation leads to the transcriptional regulation of a wide array of IFN-stimulated genes (ISG) that ordinarily protect cells from infection. Although IFNs were discovered owing to their antiviral properties, they are currently approved for treating a variety of diseases including hepatitis, multiple sclerosis, and melanoma ^10^.

Recently, STING has emerged as a regulator of nociception ^11–15^. Mice lacking STING developed mechanical allodynia, and STING agonists produce anti-nociceptive effects in neuropathic pain models^11^. Despite the observed analgesic properties of STING agonists, the role of STING in pathological pain and the mechanisms by which STING modulates and reprograms nociceptors in inflammatory pain remains unclear. Here, we found that inflammation induced by Complete Freund Adjuvant (CFA) leads to an upregulation of STING in Nav1.8+ and TRPV1+ neurons. We show that mice expressing a TRPV1 nociceptor-specific gain-of-function mutation in human STING exhibited an IFN gene signature associated with reduced heat hyperalgesia. Notably, we identified several IFN-regulated genes (IRGs) as nociceptor-specific ion channels, including TRPV1 and KChIP1, which control nociceptor excitability and hyperalgesia. Overall, our findings suggest that STING serves as a marker of nociceptor sensitization, and IRGs play a crucial role in regulating persistent pain conditions. This work provides novel mechanistic insights into the role of the STING-IFN-I pathway in the regulation of key ion channels and ion channel-associated proteins for pain resolution.

## Results

### Nociceptor STING is upregulated in response to inflammation

To study the mechanisms of resolution of inflammatory pain, we used the well-established inflammatory pain model of Complete Freund’s Adjuvant (CFA), which induces local inflammatory responses to the injection of heat-killed Mycobacterium tuberculosis. Mice developed thermal hyperalgesia by day one post CFA injection and recovered within two weeks (**Fig. 1A**), suggesting a high level of plasticity in the afferent pain pathway, particularly unmyelinated small-diameter nociceptors that undergo significant molecular and functional changes during inflammation ^16,17^. To explore how nociceptors respond to the inflammatory milieu, we used Nav1.8 Tg-TdTomato mice which mark neurons that express the voltage-gated sodium channel Nav1.8 (Scn10A)^18^. Following intraplantar CFA injection, we FACS sorted Nav1.8+ neurons (**Supp Fig. 1A**) from paw-innervating ipsilateral and contralateral lumbar (L4-L6) DRGs (**Fig. 1B**). Transcriptional analysis identified two differentially expressed genes, between ipsilateral and contralateral sides: Stimulator of Interferon Genes (STING) (Tmem173 gene) and Angiopoietin-like protein 2 (Angptl2) (**Fig. 1C**). To narrow down our analysis to TRPV1+ nociceptors that play a central role in inflammatory hyperalgesia ^19^, we repeated the experiment using a TRPV1-pHluorin mouse previously characterized ^20^ (**Supp Fig. 1B**, **Fig. 1D**). As observed with the larger population of Nav1.8+ neurons, we found that CFA inflammation enhanced STING expression in ipsilateral TRPV1+ neurons (**Fig. 1E**). Analyzing a single cell RNA sequencing dataset of mouse DRG neurons ^21^, we found that STING was co-expressed with peptidergic nociceptor-associated transcripts such as Scn10a (Nav1.8), Trpv1, Calca (CGRP), Tac1 (Substance P) and GRFα3 (**Supp Fig. 2A**). We thus performed immunostaining to determine the level of STING protein in the different subpopulations of DRG neurons: TRPV1 (PEP1.1, PEP1.2, PEP1.3, PEP1.4, NP3) and IB4 (NP1, NP2, NP3, TH)) (Supp **Fig. 2B**). Immunostaining analysis confirmed that STING was enriched in TRPV1+ peptidergic neurons (PEP1.1, PEP1.2, PEP1.3, PEP1.4, NP3) compared to IB4+ non peptidergic (NP1, NP2, NP3, TH) neurons (**Supp Fig. 2C**). Furthermore, we evaluated the expression of STING in human DRGs (**Supp Fig. 2D**). We found a high degree of colocalization between STING and Nav1.8 transcripts, confirming that STING was enriched in neuronal rather than non-neuronal cells of human DRGs (**Supp Fig. 2E**). Collectively, our results confirmed that STING is expressed in both human and mouse nociceptors.

**Figure 1.**
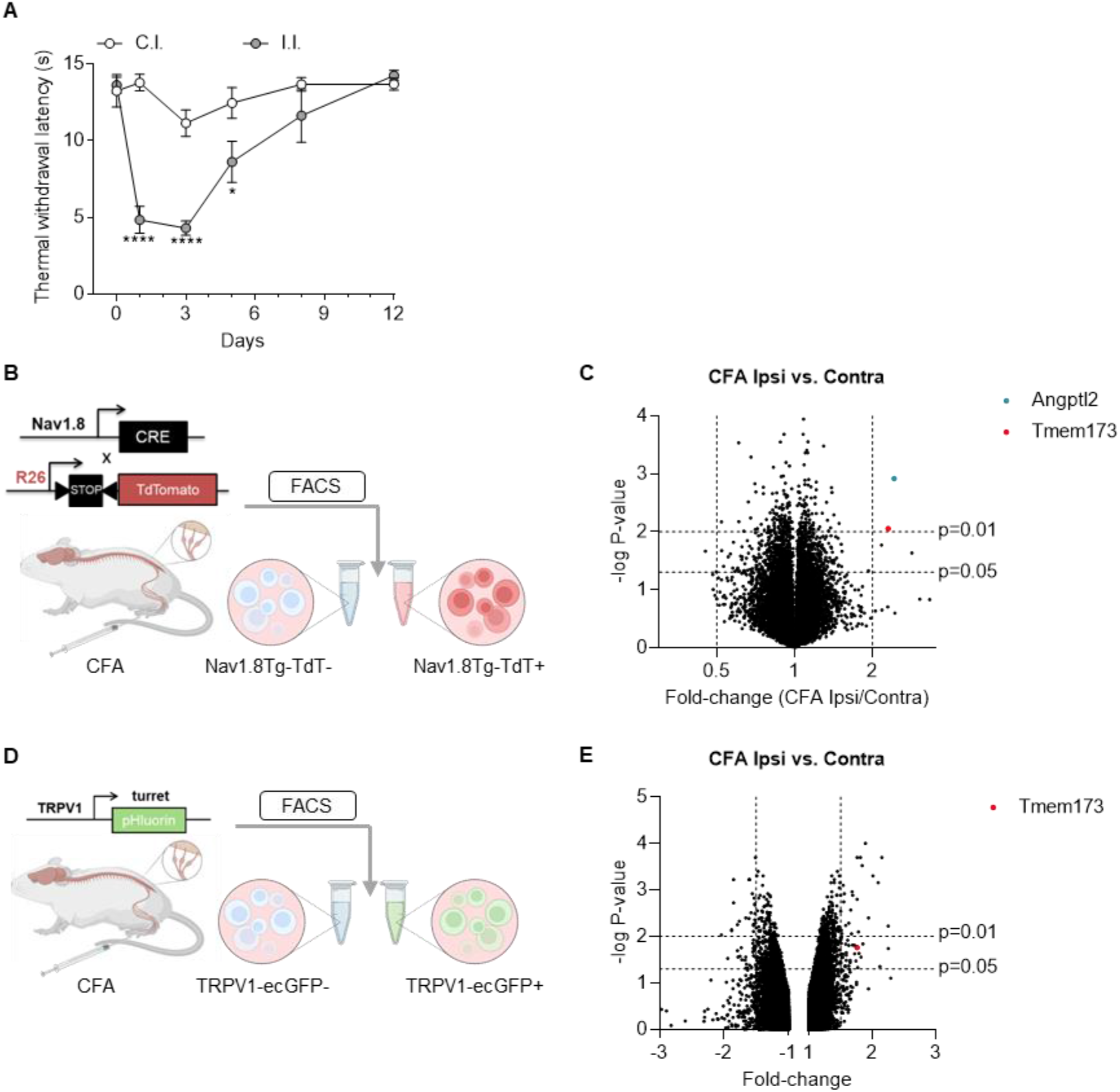
Transcriptional changes in Nav1.8 and TRPV1 neurons after CFA inflammation. **(A)** Measurement of thermal withdrawal latency in the ipsilateral (i.l.) and contralateral (c.l.) hind paws of C57BL/6J mice treated with CFA (n = 6). Statistical analysis was performed using Two-way ANOVA followed by Sidak post-hoc test (*** p < 0.001). **(B)** Experimental approach used to isolate Nav1.8Tg-TdTomato neurons for microarray analysis, 24 hours after intraplantar CFA. **(C)** Volcano plot showing transcriptional changes induced by CFA inflammation in Nav1.8Tg-TdT neurons (n = 3 mice). P-value line cut-off is p<0.01, and fold change of 2. Select transcripts of interest are highlighted in distinct colors (legend). (**D**) Experimental approach used to isolate TRPV1-pHluorin neurons for microarray analysis, 72 hours after intraplantar CFA. (**E**) Volcano plot showing transcriptional changes induced by CFA inflammation in TRPV1 neurons. P-value line cut-off is p<0.05, and fold change of 1.5. Select transcript of interest is highlighted in red (legend). Results indicate the mean ± SEM.

**Figure 2:**
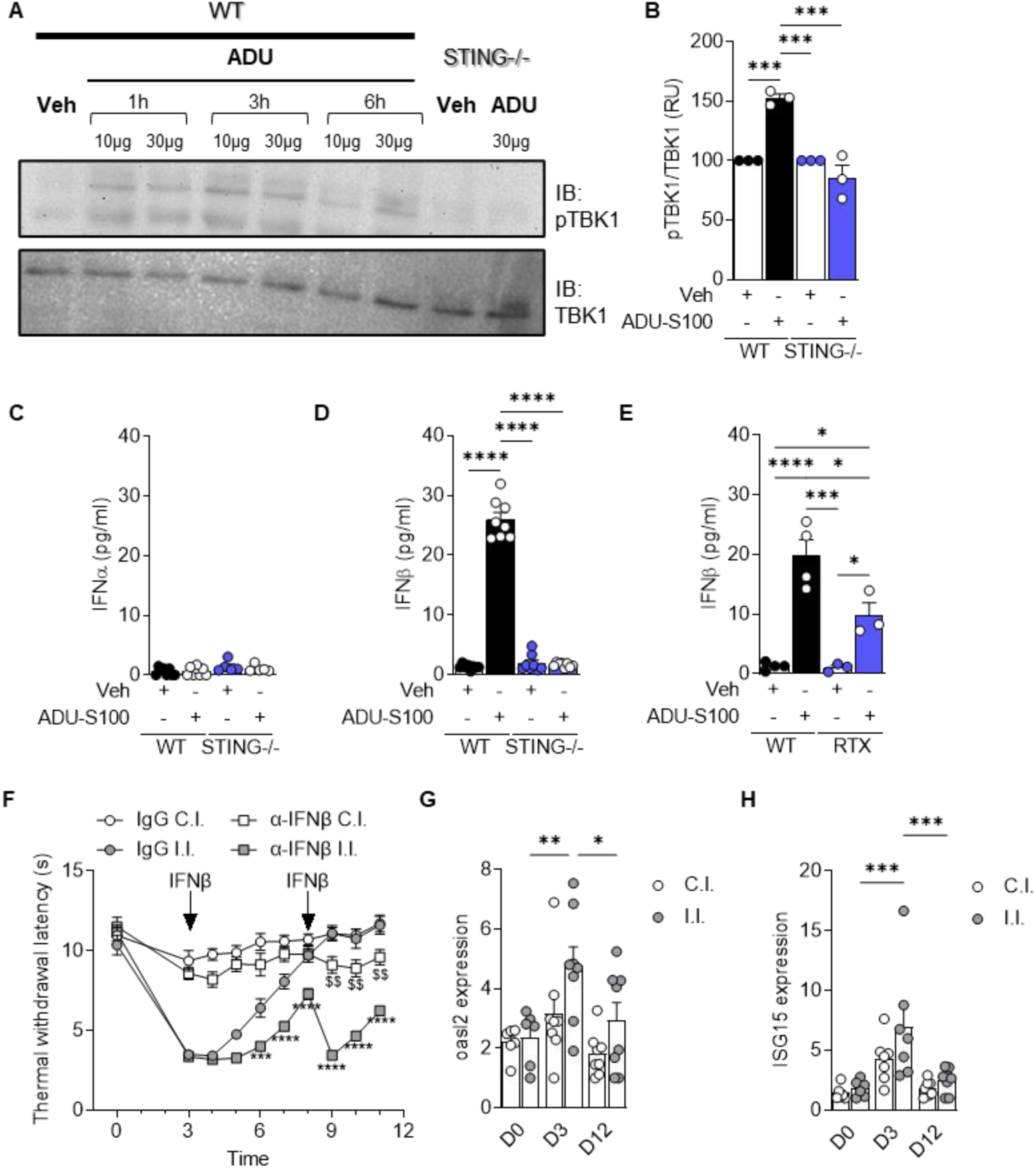
Neuronal Type I Interferon promote resolution of inflammatory pain. **(A)** Phospho-TBK1 (p-TBK1) protein level was determined by Western blot in the lumbar DRG neurons culture (L4-L6) treated with Vehicle (n = 3) or ADU-S100 (10µg or 30µg) for 1, 3 or 6 hours from WT and STING-/-mice. (**B**) Bar graph representing p-TBK1 quantification at 3 hours in response to 10µg of DU-S100. Data are normalized to TBK1 signal. Statistical analysis was performed using One-way ANOVA followed by Tukey post-hoc rest (*** p < 0.001). (**C**) IFNα and IFNβ (**D**) levels were determined by Luminex technology in the DRG culture of WT (n = 4) and STING-/- (n =4-8) DRG cultures treated with Vehicle or ADU-S100. Statistical analysis was performed using One-way ANOVA followed by Tukey post hoc test (**** p < 0.0001). (**E**) IFNβ levels were determined by Luminex technology in Vehicle (n = 4) and RTX (n = 3) pre-treated DRG neurons, stimulated with ADU-S100. Statistical analysis was performed using One-way ANOVA followed by Tukey post hoc test (* p < 0.05; *** p < 0.001; **** p < 0.0001). (**F**) Measurement of thermal withdrawal latency in the hind paws of CFA-treated C57BL/6J mice that received either an IgG control (n = 11) or an IFNβ neutralizing antibody at day 3 and day 8 post CFA injection (n = 11). Statistical analysis was performed using Two-way ANOVA followed by Tukey post-hoc test (*** p < 0.001, **** p < 0.0001 vs IgG I.l.; $$ p < 0.01 vs IgG C.l.). (**G**) OASL2 expression in both ipsilateral and contralateral lumbar DRG neurons of naïve C57BL/6J (D0, n = 6), 3 days post-CFA injection (D3, n = 8) and 12 days post-injection (D12, n = 8) mice. Statistical analysis was performed using Two-way ANOVA followed by Tukey post-hoc test (* p < 0.05,** p < 0.01). (**H**) ISG15 expression in both ipsilateral and contralateral lumbar DRG neurons of naïve C57BL/6J (D0, n = 6), 3 days post-CFA injection (D3, n = 7) and 12 days post-injection (D12, n = 8) mice. Statistical analysis was performed using Two-way ANOVA followed by Tukey post-hoc test (*** p < 0.001). Results indicate the mean ± SEM.

### Type I Interferons (IFN-I) promote resolution of inflammatory pain

STING activation in immune or epithelial cells leads to the production of Type I interferons (IFN-I) such as interferon α (IFNα) and interferon β (IFNβ) isoforms. This IFN-I response is mediated by the recruitment of TANK-binding kinase 1 (TBK1) and its phosphorylation^22^. Accordingly, activation of STING by ADU-S100 in cultured DRG neurons, led to the phosphorylation of TBK1 in WT but not STING-/- neurons (**Fig. 2A-B**). Additionally, ADU-S100 stimulation induced the production of IFNβ, but not IFNα, and this IFN-I response was dependent on the presence of STING (**Fig. 2C-D**). Thirteen distinct cell clusters have been identified in the DRG ^23^, including macrophages, fibroblasts and satellite glial cells, which express STING (http://mousebrain.org/)^21^. To assess the proportion of IFN-β produced by TRPV1-expressing neurons among all DRG cells, cultures were treated with Resiniferatoxin (RTX) to ablate TRPV1+ neurons prior to ADU-S100 exposure. RTX pre-treatment reduced the amount of IFNβ produced in response to the STING agonist by 50%, indicating that a substantial amount of IFN-I originated from TRPV1+ neurons (**Fig. 2E**).

To investigate whether IFNβ mediated the resolution of thermal hyperalgesia, we used an αIFNβ neutralizing antibody. Mice subjected to CFA received intrathecal injections of αIFNβ antibody or an IgG control on the third-and eighth-days following CFA injection. Neutralizing IFNβ delayed the resolution of thermal hyperalgesia in CFA-treated mice (**Fig. 2F**), suggesting a role of IFN-I for pain resolution. IFN-I bind to IFNα/β receptor (IFNAR) on producer and nearby cells. This autocrine and paracrine stimulation leads to the transcriptional regulation of a wide array of IFN-stimulated genes (ISG) that ordinarily protect cells from infection. We thus assessed the expression of classical ISGs, specifically OASL2 and ISG15, over time in CFA-treated mice. Our findings revealed an increase of both OASL2 and ISG15 three days after CFA injection in ipsilateral DRGs, returning to baseline levels by day 12 (**Fig. 2G-H**). These results indicate that the resolution of CFA-induced hyperalgesia is associated with a robust IFN signature.

### IFN-I/IFNAR1 signaling axis regulates thermal hyperalgesia

To investigate the contribution of neuronal IFN-I in regulating nociceptive behaviors, we expressed a STING gain of function (GOF) mutant in TRPV1 neurons. The hSTING mutation (N154S) has been reported in STING-associated vasculopathy with onset in infancy (SAVI), a type I interferonopathy associated with constitutive activity of STING ^24^. We generated TRPV1^cre^-GOF cKI mice by crossing hSTING-N154S mice with TRPV1-Cre animals (**Fig. 3A**). When assessing the efficiency of Cre recombination and the expression of hSTING-N154S in TRPV1+ neurons (**Supp Fig. 3A**), we made two important observations: 1) the number of TRPV1+ neurons in the TRPV1^cre^-GOF was reduced compared to littermate controls and 2) the large majority of TRPV1+ neurons expressed the hSTING-N154S mutation (**Supp Fig. 3B-C**). We observed a similar trend with the peptidergic GFRα3 marker, while the expression of other neuronal markers, such as SP or GFRα2, remained similar in both TRPV1^cre^-GOF and littermate neurons (**Supp Fig.4 A-F**).

**Figure 3.**
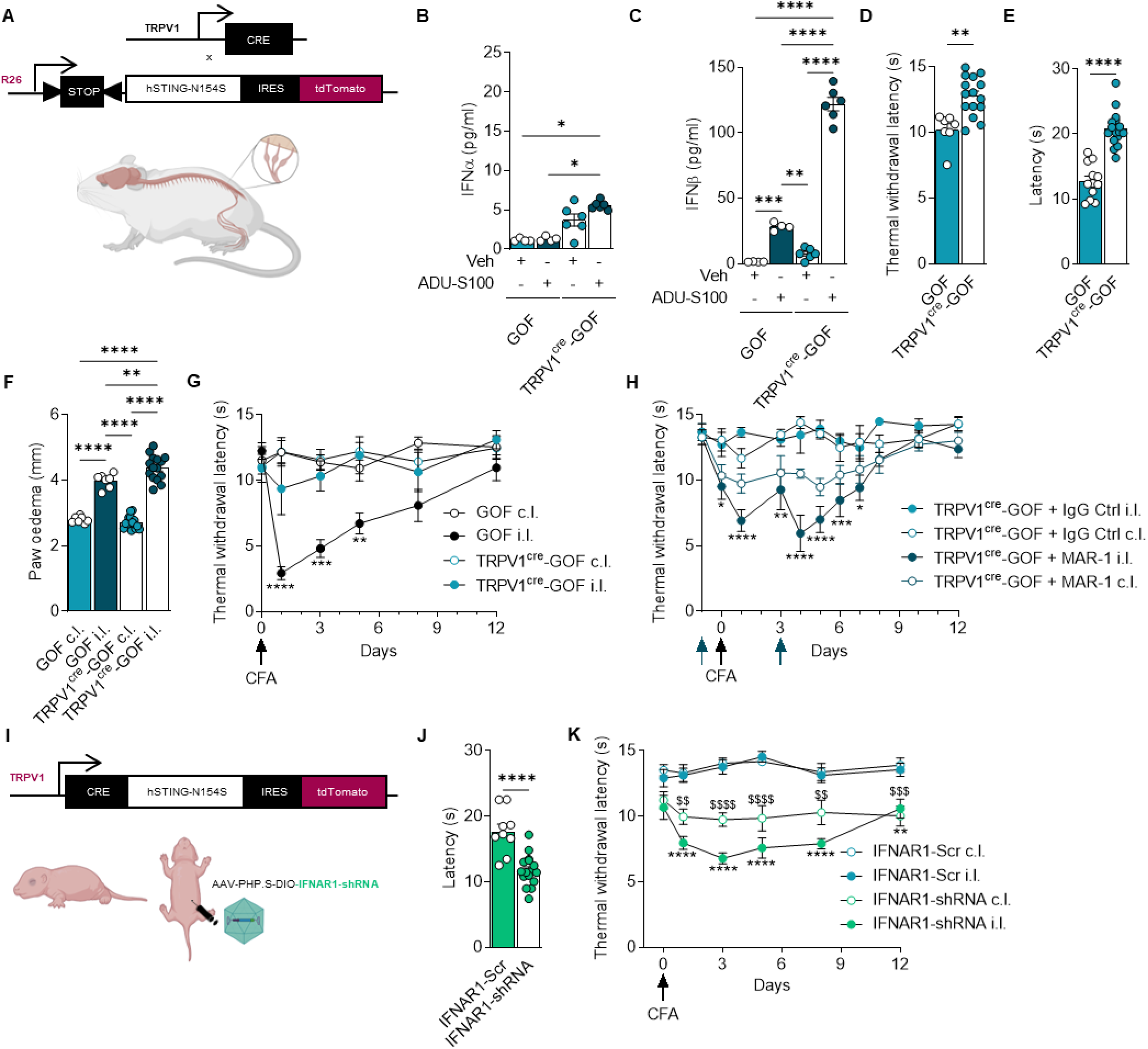
Nociceptor-specific STING-N154S gain of function reduces thermal sensitivity and heat hyperalgesia in an IFNAR1 dependent manner. **(A)** Schematic representation of the transgenic TRPV1^cre^-GOF cKi mouse design. (**B**) IFNα levels were determined by Luminex technology in DRG cultures of GOF (n = 4) and TRPV1^cre^-GOF (n = 6) mice and stimulated with ADU-S100. Statistical analysis was performed using Kruskal-Wallis followed by Dunn’s post hoc test (* p < 0.05). (**C**) IFNβ levels were determined by Luminex technology in DRG cultures of GOF (n = 4) and TRPV1^cre^-GOF (n = 6) mice and stimulated with ADU-S100. Statistical analysis was performed using One-Way ANOVA by Tukey post hoc test (** p < 0.01; *** p < 0.001; **** p < 0.0001). (**D-E**) Measurement of thermal sensitivity in hind paws of naïve TRPV1^cre^-GOF (n = 15-16) and littermate control GOF mice (n = 7-12) using the Hot plate (**D**) or Hargreaves test (**E**). Statistical analysis was performed using t-test (**D**) or Mann-Whitney test (**E**) (** p < 0.01, **** p < 0.0001). (**F**) Measurement of edema in the ipsilateral and contralateral hind paws of CFA-treated GOF mice (n = 9) and TRPV1cre-GOF mice (n = 8). Statistical analysis was performed using One-Way ANOVA followed by Tukey post-hoc test (** p < 0.01, **** p < 0.0001). (**G**) Measurement of thermal withdrawal latency in the ipsilateral and contralateral hind paws of GOF (n = 9) and TRPV1^cre^-GOF (n = 8) mice that were treated with CFA. Statistical analysis was performed using Two-way ANOVA followed by Tukey post-hoc test (* p < 0.05, ** p < 0.01, *** p < 0.001). (**H**) Measurement of thermal withdrawal latency in the hind paws of CFA-treated TRPV1^cre^-GOF mice that received either an IgG control (n = 5) or IFNAR1 neutralizing antibody (MAR-1) prior to CFA injection and at day three post CFA injection (n = 6). Statistical analysis was performed using Two-way ANOVA followed by Tukey post-hoc test (* p < 0.05, ** p < 0.01, *** p < 0.001, **** p < 0.0001). (**I**) Newly born pups (postnatal day 5) were administered 10 µL of AAV-PHP.S-DIO-IFNAR1-shRNA or AAV-PHP.S-DIO-scrambled-shRNA intraperitoneally. (**J**) Measurement of thermal sensitivity in the hind paws of AAV-DIO-scram-shRNA (n = 9) and AAV-DIO-IFNAR1-shRNA (n = 16) injected mice. Statistical analysis was performed using t-test (**** p < 0.0001). (**K**) Measurement of thermal withdrawal latency in the ipsilateral and contralateral hind paws of AAV-DIO-scram-shRNA (n = 7) and AAV-DIO-IFNAR1-shRNA (n = 9) infected mice treated with CFA. Statistical analysis was performed using Two-way ANOVA followed by Tukey post-hoc test (** p < 0.01, **** p < 0.0001 vs AAV-DIO-scram-shRNA i.l.; ($$ p < 0.01, $$$$ p < 0.0001, vs AAV-DIO-scram-shRNA c.l.). Results indicate the mean ± SEM.

**Figure 4.**
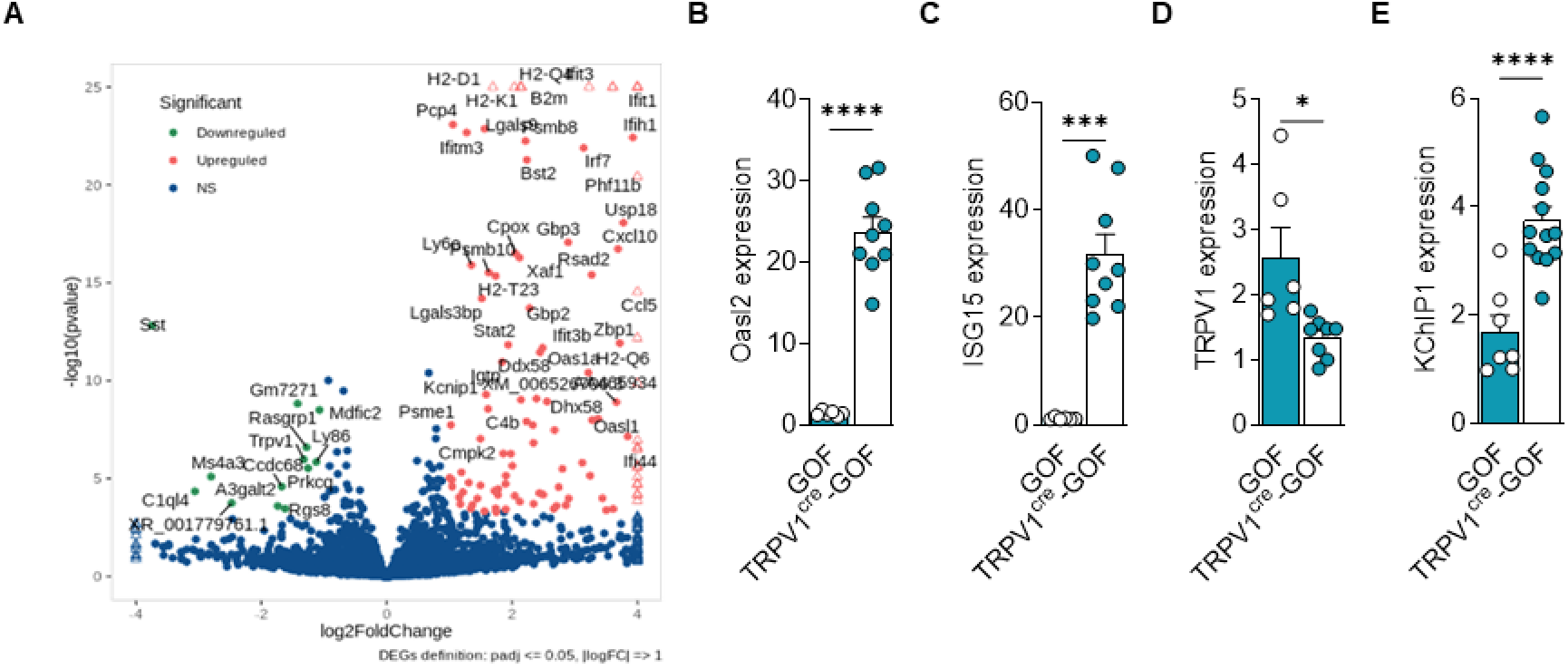
Nociceptor-specific STING-N154S gain of function induces IFN-I production and expression of Interferons Stimulated Genes (ISGs) in DRGs. (**A**) Volcano plot representation of genes regulated in naive TRPV1^cre^-GOF cKi mice. Genes that pass a threshold of log1.5-fold change in differential expression analysis are colored in green when they are downregulated and red when they are upregulated. (**B**) OASL2 expression in lumbar DRG neurons of naïve GOF (n = 6) and TRPV1^cre^-GOF (n = 9) mice. Statistical analysis was performed using t-test (**** p < 0.0001). (**C**) ISG15 expression in lumbar DRG neurons of naive GOF (n = 6) and TRPV1^cre^-GOF (n = 9) mice. Statistical analysis was performed using Mann-Whitney test (*** p < 0.001). (**D**) TRPV1 expression in lumbar DRG neurons of naive GOF (n = 6) and TRPV1^cre^-GOF (n = 8) mice. (**E**) KChIP1 expression in lumbar DRG neurons of naive GOF (n = 7) and TRPV1^cre^-GOF (n = 13) mice. Statistical analysis was performed using t-test (**** p < 0.0001). Statistical analysis was performed using t-test (* p < 0.05). Results indicate the mean ± SEM.

We next examined the expression of TdTomato in the central terminals of nociceptors in the superficial dorsal horn of the spinal cord (**Supp Fig. 5**). Immunohistochemical analysis led to two observations: 1) hSTING-N154S expression occurs mainly in Substance P (peptidergic) (**Supp Fig. 5A**); and GFRα3 (peptidergic) nociceptors (**Supp Fig. 5B**), with fewer IB4 (non-peptidergic) nociceptors (**Supp Fig. 5C**); and 2) the neuroanatomical organization of peptidergic and non-peptidergic fibers was conserved in TRPV1^cre^-GOF mice.

**Figure 5.**
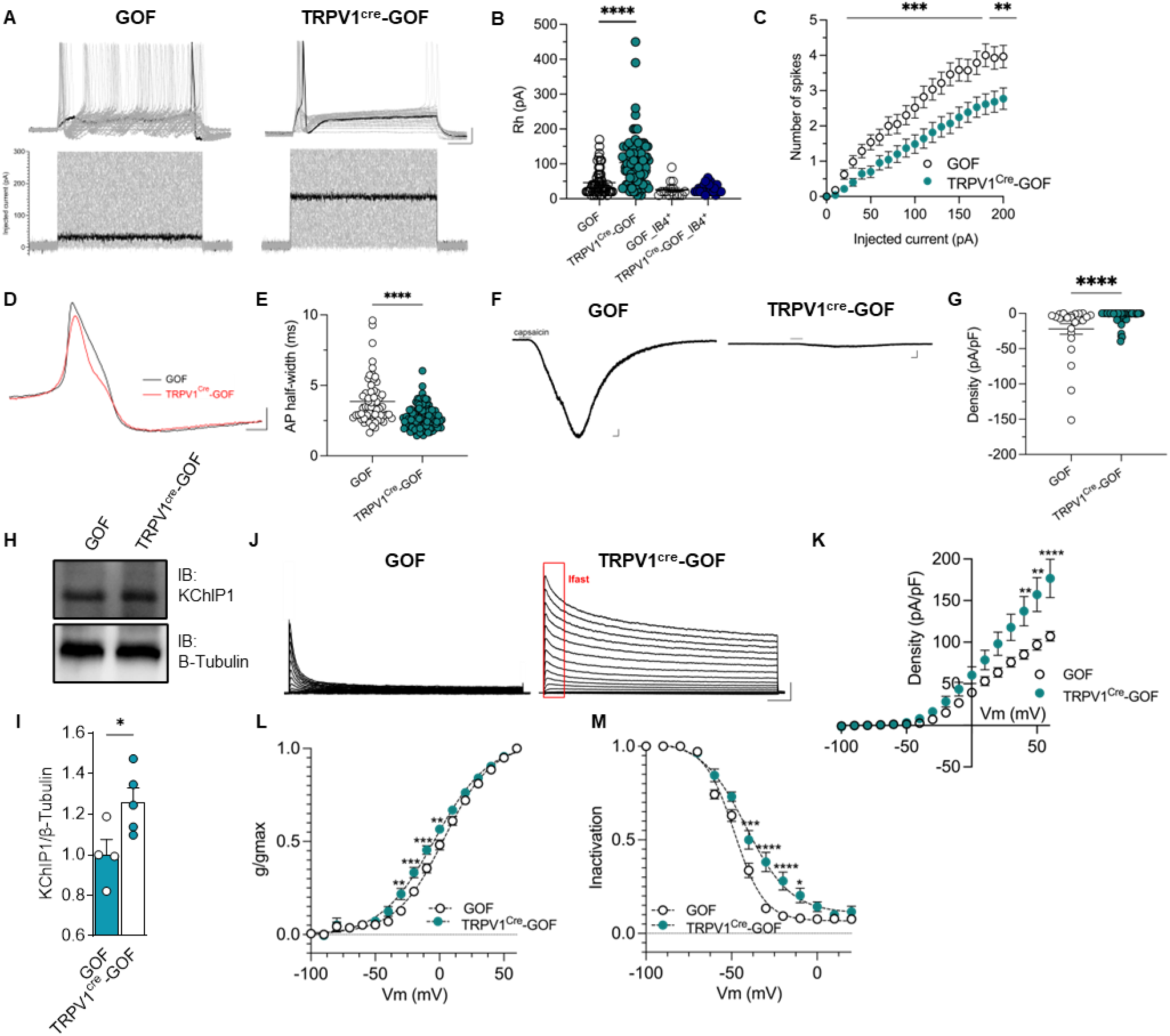
ISGs alter nociceptor properties through TRPV1 downregulation and KChIP1 expression. (**A**) Representative current clamp recording of evoked action potentials recorded in TRPV1 neurons (TdTomato+) from GOF (left) and TRPV1^cre^-GOF (right) mice. Cells were injected with a 500 ms current pulse with an increment of 10 pA and an interval of 5 seconds. Scale bars: 20 mV/50 ms. **(B)** Rheobase data recorded in TRPV1 (TdTomato+) and non-peptidergic (IB4+) neurons from GOF mice (45.33 ± 4.72 pA, n=61 TRPV1+ neurons; 25.63 ± 5.40 pA, n=16 IB4+ neurons) and TRPV1^cre^-GOF mice (103.70 ± 6.85 pA, n=101 TRPV1+ neurons; 29.09 ± 2.86 pA, n=22 IB4+ neurons). Statistical analysis was performed using the Kruskal-Wallis test with Dunn’s multiple comparison test. **** p < 0.0001. **(C)** Number of spikes as a function of injected current in TRPV1 neurons from TRPV1^cre^-GOF mice and GOF littermate controls. Statistical analysis was performed using a two-way ANOVA test with Tukey’s post-hoc test. (** p < 0.01, *** p < 0.001). **(D)** Representative action potentials recorded in TRPV1 neurons from GOF (black) and TRPV1^cre^-GOF (red) mice. Scale bars: 20 mV/50 ms. **(E)** Action potential half-width recorded in TRPV1 neurons from GOF (3.86 ± 0.21 ms, n=61) and TRPV1^cre^-GOF (2.73 ± 0.82 ms, n=101) mice. Statistical analysis was performed using the Mann-Whitney test. **** p < 0.0001. **(F)** Representative currents induced by 100 nM of capsaicin in TRPV1 neurons from GOF (left) and TRPV1^cre^-GOF (right) mice. Scale bars: 200 pA/10 s. **(G)** Current density evoked by capsaicin in TRPV1 neurons from GOF (-21.93 ± 7.51 pA/pF, n=25) and TRPV1^cre^-GOF (-2.85 ± 0.96 pA/pF, n=70) mice. Statistical analysis was performed using the Mann-Whitney test. ****p<0.0001. (**H**) Western blot of KChIP1 and β-tubulin from lumbar DRG lysates. **(I)** Quantification of KChIP1 protein level in DRG lysates from naïve GOF (n = 4) and TRPV1^cre^-GOF (n = 5) mice, measured by Western blot. Statistical analysis was performed using the Mann-Whitney test (* p < 0.05). **(J)** Representative outward potassium currents recorded in TRPV1 neurons from GOF (left traces) and TRPV1^cre^-GOF (right traces) mice. Scale bars: 1 nA/100 ms. **(K)** Average current-voltage relationship from neurons recorded in (J). GOF (n=10) and TRPV1^cre^-GOF (n=17). **(L)** Steady-state activation and inactivation **(M)** from neurons recorded in (J). Activation; GOF: V1/2 = 3.567 mV, K=18.68 mV, n=17; TRPV1^cre^-GOF: V1/2 = -3.779 mV, K=21.20 mV, n=17. Inactivation; GOF: V1/2 = -48.31 mV, K=-8.72 mV, n=18; TRPV1^cre^-GOF: V1/2 = -41.56 mV, K=-13.01 mV, n=25. All steady-state plots were fitted with Boltzmann functions to derive V1/2 and K values. Statistical analysis was performed using a two-way ANOVA test with Tukey’s post-hoc test. (** p < 0.01, *** p < 0.001, **** p < 0.0001). Results indicate the mean ± SEM.

Next, we assessed IFN-I production in TRPV1^cre^-GOF animals. Cultured DRG neurons from TRPV1^cre^-GOF mice exhibited an elevated level of type I IFNα and β in the absence of STING stimulation. In addition, ADU-S100 induced a >ten-fold increase in IFNβ, but not IFNα, compared to untreated neurons (**Fig. 3B-C**), and ADU-induced increase of both α and β IFN was six-fold larger in cKI animals versus littermate controls (**Fig. 3B-C**).

We then wanted to determine whether constitutive IFN-I signalling could affect sensory or pain-like behaviors. While baseline mechanical sensitivity was not altered in naïve TRPV1^cre^-GOF mice (**Supp Fig. 6A)**, thermal sensitivity as measured by both the Hot plate and Hargreaves test was lower than littermate controls (**Fig. 3D-E**). In addition, stereotypical mouse behaviors such as climbing, rearing, eating, drinking, grooming, scratching and distance moved were normal in TRPV1^cre^-GOF cKI animals (**Suppl Fig. 6C-J**), indicating that IFN-I production by TRPV1+ neurons mainly controlled thermal nociception.

**Figure 6.**
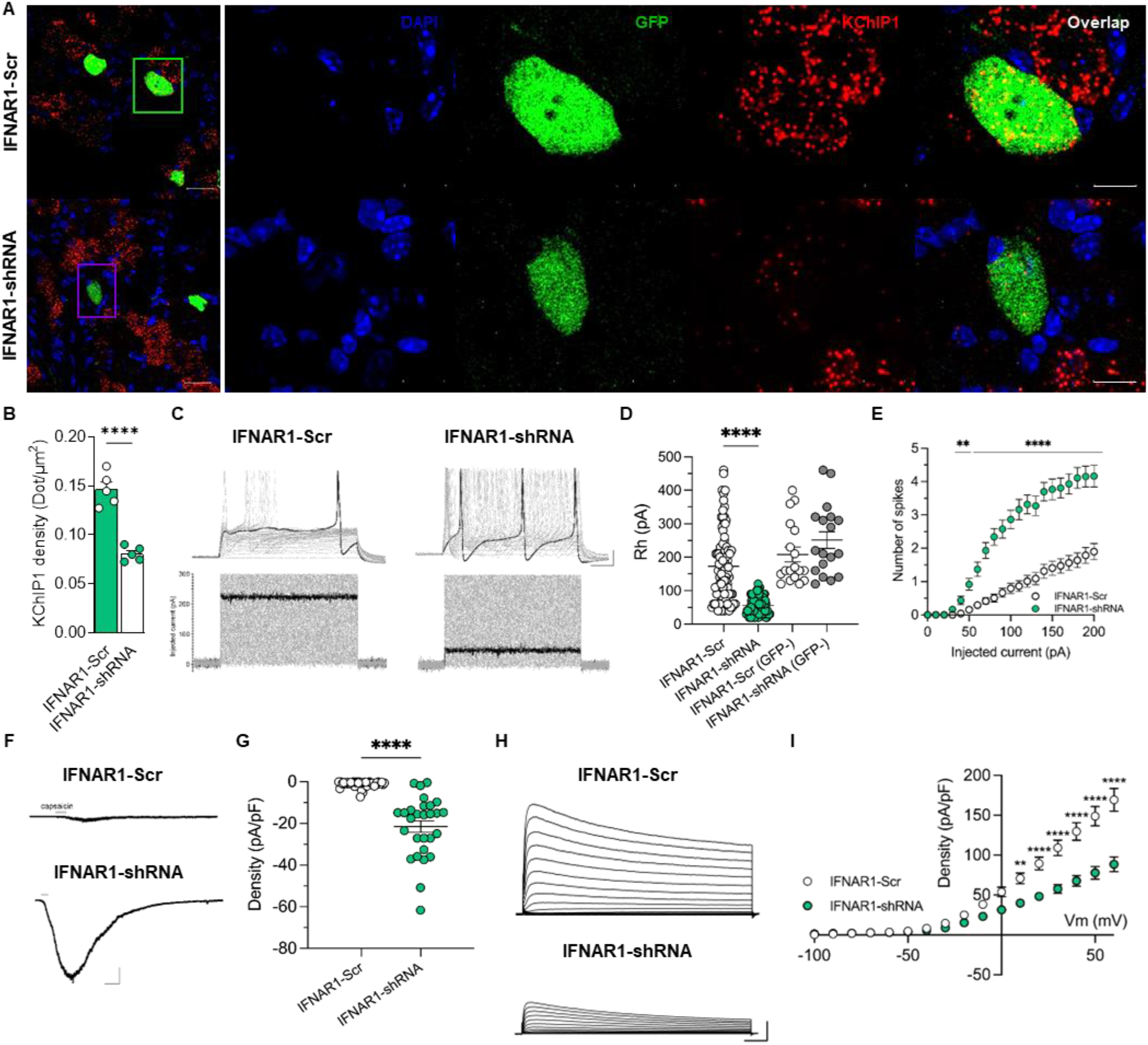
IFNAR1 depletion in nociceptors restores electrophysiological properties. (**A**) Representative confocal image of TRPV1 neurons from TRPV1^Cre^-GOF mice injected with AAV-DIO-Scr-shRNA (n = 5) or AAV-DIO-IFNAR1-shRNA (n = 5). Images represent DAPI staining, AAV-GFP expression (green) and KChIP1 transcripts (red) by RNAscope. Scale bars: 25 µm and 10 µm on the cropped images. **(B)** Quantification of KChIP1 density measured by the number of transcripts represented by dots per surface unit in AAV-infected TRPV1 neuron. Statistical analysis was performed using unpaired t-test (**** p < 0.0001). **(C)** Representative current clamp recording of TdTomato+/GFP+ TRPV1 neurons from TRPV1^Cre^-GOF mice infected with DIO-Scram-shRNA (left) or DIO-IFNAR1-shRNA (green) AAVs. Cells were injected with 500 ms current pulses with an increment of 10 pA and an interval of 5 seconds. Scale bars: 20 mV/50 ms. **(D)** Measurement of Rheobase in infected (GFP+) and non-infected (GFP-) TRPV1 neurons (TdTomato+) recorded in (D). Data are presented as dot plots with mean values (172.8 ± 11.04 pA, n=89 for IFNAR1-Scr neurons; 56.13 ± 2.75 pA, n=80 for IFNAR1-shRNA infected neurons; and 208.30 ± 21.89 pA, n=18 for IFNAR1-Scr neurons; 251.10 ± 24.77 pA, n=18 for IFNAR1-shRNA non-infected neurons). Statistical analysis was performed using the Mann-Whitney test. ****p<0.0001. **(E)** Number of spikes evoked by injected current in TRPV1 (TdTomato+) and AAV infected (GFP+) neurons. Statistical analysis was performed using a two-way ANOVA test with Tukey’s post-hoc test. (** p < 0.01, **** p <0.0001). **(F)** Representative capsaicin-evoked current in TdTomato+/GFP+ TRPV1 neurons from TRPV1^Cre^-GOF mice infected with IFNAR1-Scr (left) and IFNAR1-shRNA (right) AAVs. Scale bars: 200 pA/20 s. **(G)** Current density evoked by capsaicin in cells represented in G); IFNAR1-Scr (-1.13 ± 0.23 pA/pF, n=41) and IFNAR1-shRNA (-21.56 pA/pF, n=28) neurons. Statistical analysis was performed using the Mann-Whitney test. **** p < 0.0001. (**H**) Representative outward potassium currents recorded in TRPV1 neurons from TRPV1^Cre^-GOF infected with IFNAR1-Scr (left traces) and IFNAR1-shRNA (right traces) AAVs. Scale bars: 1 nA/100 ms. **(I)** Average current-voltage relationship in TRPV1 neurons from TRPV1^Cre^-GOF mice infected with IFNAR1-Scr (n=31) or IFNAR1-shRNA AAVs (n = 25). Statistical analysis was performed using a two-way ANOVA test with Tukey’s post-hoc test. (** p < 0.01, **** p < 0.0001). Results indicate the mean ± SEM.

We next assessed the thermal hyperalgesia of TRPV1^cre^-GOF mice after CFA. Although a slightly more pronounced inflammatory response was observed in the ipsilateral paw of TRPV1^cre^-GOF mice compared to littermate controls (**Fig. 3F**), TRPV1^cre^-GOF mice exhibited negligible inflammatory thermal hyperalgesia (**Fig. 3G**). However, mechanical allodynia remained unaffected (**Supp Fig. 6B)**. This analgesic effect was not due to a systemic release of IFN-I or a change in pro-or anti-inflammatory cytokines (**Supp Fig. 7A-F**, **Table S1**). To assess if the type I IFN receptor (IFNAR1) mediated the anti-hyperalgesic effect of the GOF, we used an αIFNAR1 blocking antibody (MAR1). TRPV1^cre^-GOF mice received intrathecal injections of MAR1 antibody (1 µg in 10 µl) or IgG control (1µg in 10 µl) prior to, and three days following CFA injection. Inhibition of IFNAR1 by MAR1 antibody restored thermal hyperalgesia in CFA-treated TRPV1^cre^-GOF mice (**Fig. 3H**). Because IFNAR1 is expressed in both neuronal and non-neuronal cells of the DRG ^11,21^, we assessed the specific role of IFNAR1 expressed in TRPV1+ fibers by deleting IFNAR1 specifically in nociceptors. TRPV1^cre^-GOF neonates (P5) received either an i.p. injection of AAV expressing Cre-inducible IFNAR1-shRNA or Scramble control (**Fig. 3I**). At six weeks, we confirmed that neurons expressing the STING-GOF mutant were infected with the viral construct (**Supp Fig. 8A**). Downregulation of IFNAR1 transcripts was demonstrated by RNAscope, validating the efficacy of the IFNAR1-shRNA vector (**Supp Fig. 8B**). Depletion of IFNAR1 in TRPV1^cre^-GOF neurons restored normal thermal sensitivity (**Fig. 3J**). Importantly, mice treated with IFNAR1-shRNA displayed thermal hyperalgesia compared to IFNAR1-Scr mice after CFA injection (**Fig. 3K**). Overall, these results show that activation of neuronal STING might promote resolution of pain after inflammation, at least in part via autocrine type I IFN-IFNAR1 signaling.

**Figure 7.**
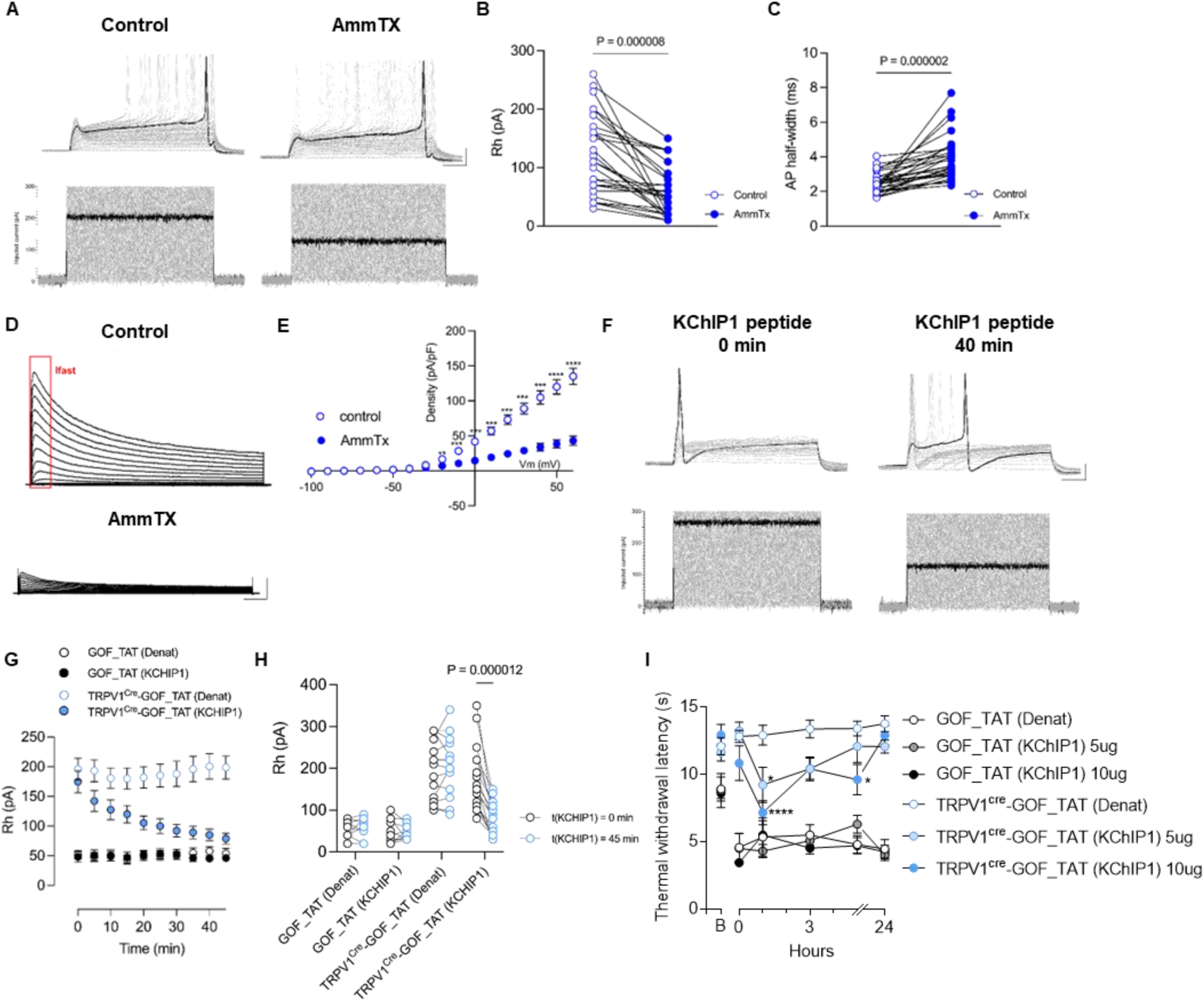
KChIP1/Kv4 interaction promotes the antinociceptive effect of ISGs. **(A)** Representative current clamp recording of TRPV1 neurons from TRPV1^cre^-GOF mice, in control condition (left traces) and after AmmTx3 application (right traces). Cells were injected with 500 ms current pulses with an increment of 10 pA and an interval of 5 seconds. Scale bars: 20 mV/50 ms. **(B)** Measurement of the change in Rheobase (left graph) and (**C**) action potential half-width (right graph) induced by AmmTx3 application in TRPV1 neurons from TRPV1^cre^-GOF mice (n=29 neurons). Statistical analysis was performed using a paired t-test. **(D)** Representative outward potassium currents recorded in TRPV1 neurons from TRPV1^cre^-GOF mice, in control condition (left traces) and after AmmTx3 application (right traces). Scale bars: 1 nA/100 ms. **(E)** Average current-voltage relationship obtained from the cells recorded in D (n=9). **(F)** Representative current clamp recording of TRPV1 neurons, from TRPV1^cre^-GOF mice, treated with a TAT-conjugated KChIP1 peptide for 40 min. Cells were injected with 500 ms current pulses with an increment of 10 pA and an interval of 5 seconds. Scale bars: 20 mV/50 ms. Note that intra-pipette administration of KChIP1peptide reduces rheobase. **(G)** Time-dependent effect of the TAT-conjugated KChIP1, versus denaturated control peptide, on the rheobase of TRPV1^Cre^-GOF neurons (n= 15 and 17, respectively) and GOF control neurons (n=10 and 11, respectively). **(H)** Measurement of the rheobase at t=0 and 45 min. post KChIP1 or denaturated peptide, in TRPV1 neurons from TRPV1^Cre^-GOF mice (n=15 for TAT-Denat, n=17 for TAT-KChIP1); or GOF littermate controls (n=10 for TAT-Denat, n=12 for TAT-KChIP1). Statistical analysis was performed using a paired t-test. **(I)** Measurement of thermal withdrawal latency in the hind paws of both CFA-treated GOF and TRPV1^cre^-GOF mice treated with the KChIP1 blocking peptide 5µg (n = 7;7) or 10µg (n = 6;7); or its denaturated control (n = 6;7), at day three post CFA injection. Statistical analysis was performed using Two-way ANOVA followed by Tukey post-hoc test (* p < 0.05, **** p < 0.0001). Results indicate the mean ± SEM.

**Figure 8:**
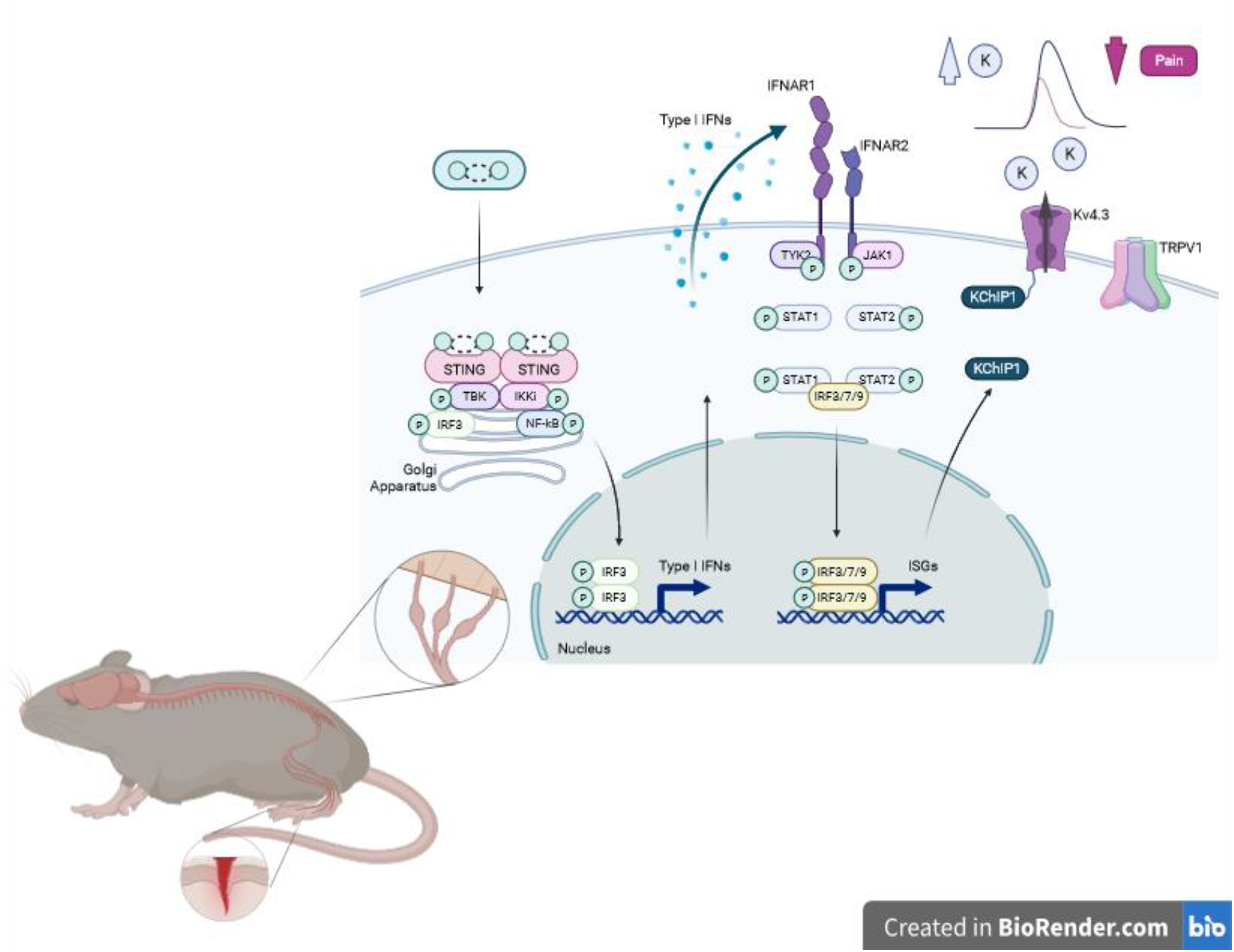
Schematic representation of pain resolving effects of the STING-IFN-I pathway in inflammatory pain models. Upregulation of nociceptor STING during inflammation stimulates TANK Binding Kinase 1 (TBK1). TBK1 phosphorylates IFN regulatory factor 3 (IRF3), which controls the production of Type I IFN, including interferon β (IFNβ). Type I interferons (IFN-I) bind to IFNα/β receptor (IFNAR) on TRPV1 nociceptors, to initiate the transcriptional regulation of hundreds of IFN-regulated genes (IRGs), which reduces heat hyperalgesia (TRPV1 downregulation) and nociceptor excitability (KChIP1 upregulation), promoting pain resolution.

### Interferon stimulated genes reduce intrinsic excitability of nociceptors

To evaluate the effect of IFN-I signaling on nociceptors, we conducted bulk RNAseq on DRG from TRPV1^cre^-GOF mice. Comparative transcriptomic profiling showed a marked interferon signature associated with a dozen of downregulated genes and >100 upregulated Interferon Stimulated Genes (ISGs) in DRGs from naïve TRPV1^cre^-GOF cKi mice, compared to GOF littermate controls (**Fig. 4A**). Among upregulated genes, we found classical ISGs known to trigger protective defense mechanisms against pathogens or tumors: OASL, IFITM3, ISG15, CCL5, USP18 mRNAs. These findings indicated that constitutive activation of STING effectively promoted ISG expression in TRPV1 nociceptors, thus validating our TRPV1^cre^-GOF cKi model to study the interferon gene responses in these neurons. While the function of ISGs enables host defense, and allows cells to recover to normal function, only a handful of these genes have been studied in detail ^10^, and none of them in the context of neuronal plasticity. Interestingly, nociceptor-specific ion channels, including TRPV1, TRPA1 and TRPC3, were downregulated in TRPV1^cre^-GOF DRGs, whereas the A-type Kv channel regulating protein (KchIP1) known to suppress excitability, was upregulated (**Fig. 4A**). Gene expression quantification by RT-qPCR confirmed the increase in OASL2 (**Fig. 4B**), ISG15 (**Fig. 4C**), and Kcnip1 (KChIP1) transcripts (**Fig. 4D**) in TRPV1^cre^-GOF neurons, whereas TRPV1 mRNAs was significantly reduced (**Fig. 4E**) corroborating the reduction of TRPV1+ neurons in the TRPV1^cre^-GOF (**Supp Fig. 3C**). To investigate the functional impact of ISGs on nociceptors, we performed electrophysiological recordings of TdTomato+ TRPV1^cre^-GOF nociceptors (**Fig. 5A**). While the AP amplitude (**Supp Fig.9A**) and the resting membrane potential (RMP) (**Supp Fig.9B**) were unchanged, an increase in Rheobase (**Fig. 5A-B**) and Rin (**Supp Fig.9C)** were measured in TRPV1^cre^-GOF neurons compared to littermate controls (GOF). In contrast, the rheobase of non-peptidergic IB4+ neurons from TRPV1^cre^-GOF mice were similar to IB4+ neurons isolated from littermate controls (**Fig. 5B** and **Supp. Fig. 10A-B**), suggesting that IFN-induced hypoexcitability was specific to TRPV1 neurons in TRPV1^cre^-GOF mice. Similarly, the number of AP evoked by depolarizing current was lower in TRPV1^cre^-GOF compared to littermate control neurons (**Fig. 5C**). Lastly, when looking at the AP half width, TRPV1^cre^-GOF mice had a shorter AP duration, indicating larger hyperpolarizing K+ currents in TRPV1^cre^-GOF neurons (**Fig. 5D-E**). Given that TRPV1 gene expression was reduced in TRPV1^cre^-GOF DRGs (**Fig. 4E** **and Supp.** Fig. 3C), we assessed capsaicin-evoked currents in isolated DRG neurons. We found a pronounced decrease in TRPV1 current density (**Fig. 5F-G**). Consistent with the observation of KChIP1 upregulation by sequencing (**Fig. 4D**), we found that KChIP1 protein level was augmented in TRPV1^cre^-GOF lumbar DRGs (**Fig. 5H-I**). As KChIP1 serves as a specific β-subunit for Kv4 channels^25,26^, we analysed the electrophysiological properties of A-type K+ currents in TRPV1^cre^-GOF neurons. Notably, the Kv4-mediated A-type current was larger in TRPV1^cre^-GOF neurons (**Fig. 5J-K**), supporting an increase in Kv4 channel opening or surface trafficking, induced by KChIP1 upregulation. Accordingly, the steady-state activation of A-type Kv4 channels was shifted to more hyperpolarized potentials (**Fig. 5L**), while the steady state inactivation was delayed in TRPV1^cre^-GOF neurons (**Fig. 5M**), indicating a regulation of Kv4 channel gating by KChIP1. Taken together, our results show that ISGs regulate the expression of ion channels and channel-interacting proteins to enhance the threshold of nociceptor activation and reduce the excitability of nociceptive neurons.

### IFNAR1 depletion normalizes ion channel expression and electrophysiological properties

To understand if the electrophysiological phenotype of TRPV1^cre^-GOF neurons was mediated by IFN-I/IFNAR1 signaling, TRPV1^cre^-GOF neonates (P5) received either an i.p. injection of Cre-inducible AAV expressing IFNAR1-shRNA or Scramble control. Depletion of IFNAR1 in TRPV1^cre^-GOF neurons restored normal KChIP1 expression (**Fig. 6A-B**). Then, DRG neurons were dissociated and electrophysiological recording of AAV-infected or non-infected TRPV1^cre^-GOF neurons was performed (**Fig. 6C**). Notably, IFNAR1-shRNA infected neurons had a much lower Rheobase compared to non-infected or IFNAR1-scr infected TRPV1^cre^-GOF neurons (**Fig. 6D**). This was consistent with an increase in evoked AP firing observed in IFNAR1-shRNA infected cells (**Fig. 6E**). In addition, IFNAR1 depletion was able to restore TRPV1 current density (**Fig. 6F-G**) and reduce Kv4-mediated A-type current (**Fig. 6H-I**) induced by IFN-I.

### KChIP1/Kv4 interaction promotes the antinociceptive effect of ISGs

Our results indicate that Kcnip1 (KChIP1) is a key interferon-regulated gene in response to nociceptor STING activation. Since we observed an increase in A-type K+ current and KChIP1 is an integral subunit component of the native Kv4 channel complex, we wondered how the Kv4-KChIP1 subunit complex could contribute to the antinociceptive effect of ISGs. To investigate this, we first tested Ammtx3, a specific Kv4 channel blocker. In TRPV1^cre^-GOF nociceptors, Ammtx3 decreased the high Rheobase, (**Fig. 7A-B**), prolonged the AP duration (**Fig. 7C**) and reduced A-type Kv current density (**Fig. 7D-E**), implying an IFN-I dependent upregulation of A-type K+ current induced by the interaction between KChIP1and Kv4.

To evaluate this hypothesis, we used a TAT-conjugated KChIP1 interfering peptide to disrupt the KChIP1-Kv4.3 or KChIP1-Kv4.1 subunit complex. Intrapipette administration of the KChIP1 peptide (20 nM) in TRPV1^cre^-GOF neurons, led to a gradual reduction of the high Rheobase, over time (**Fig. 7F-G**). The effect of KChIP1 peptide was significant after 20 min and reached saturation at 45 min post-infusion. In contrast, no time dependent inhibition of the rheobase was observed with a heat-denaturated KChIP1 peptide or in the absence of KChIP1 upregulation (GOF littermate neurons) (**Fig. 7F-H**). Having validated the efficacy of the KChIP1-Kv4 disrupting peptide, we investigated whether we could target the subunit association in TRPV1^cre^-GOF mice. Intrathecal infusion of KChIP1 peptide, at 5 and 10 µg, revealed thermal hyperalgesia for three hours, in TRPV1^cre^-GOF mice, following intraplantar CFA (**Fig. 7I**). The same peptide, administered to GOF littermates, did not disrupt CFA-induced hyperalgesia. Together, our results establish that the KChIP1-Kv4 interaction is responsible for the IFN-I-induced antinociception, downstream of neuronal STING.

## Discussion

In this study, we have identified neuronal IFN-I as a critical regulator of ion channels and channel-interacting proteins in nociceptors during the resolution of inflammation. Our findings highlighted KChIP1 and TRPV1 as downstream effectors of IFN-I, playing a pivotal role in the resolution of hyperalgesia and the restoration of normal excitability following sensitization.

We show that tissue inflammation leads to an increase in STING, a cytosolic DNA sensor, within nociceptors. Constitutive activation of STING triggers TBK1 signaling and IFNβ production, which is sufficient to induce an anti-viral ISG response, ultimately reprogramming the gene expression profiles of nociceptors. Consequently, this process results in a nociceptor specific IFNAR1 dependent reduction in excitability and inflammatory hyperalgesia (**Fig. 8**).

Our findings not only highlight STING as a valuable marker for nociceptor sensitization but also underscore the importance of IFN-regulated genes, particularly KChIP1, in the control of persistent pain conditions.

Nociceptors are specialized sensory neurons that detect a wide range of danger signals, including those from pathogens or the host, through the expression of PRRs such as STING ^8^. The cGAS-STING pathway which detects cytosolic DNA, is evolutionarily conserved among diverse animal lineages ^8,22^. Consistent with previous studies ^8,21^, we found expression of STING in nociceptors of both mouse and human dorsal root ganglion (DRG) neurons. We demonstrate that during inflammation, the expression and activation of STING were upregulated in TRPV1+ neurons, which are responsible for detecting and responding to noxious stimuli. Certain pathogens, such as SARS-CoV-2, have evolved mechanisms to suppress sensory neurons related to smell and taste, and not all viral infections exhibit symptoms. While further investigation is needed to elucidate whether this nociceptor-specific STING-IFN-I signaling contributes to antiviral responses, it may facilitate the resolution of inflammation, leading to the downregulation of pain once the viral infection has been cleared.

STING responds to both self and non-self insults. While complete Freund’s adjuvant (CFA), which contains a suspension of heat-killed mycobacteria, enhances IFN-I production, heat-inactivated Mycobacterium tuberculosis (Mtb) does not have the same effect ^27,28^. However, the adjuvant activity of CFA stems from its ability to stimulate a local innate immune response, leading to a delayed hypersensitivity reaction and an intense inflammatory response at the site of injection ^29^. Interestingly, NF-kB signaling in response to cytokines such as IL-6 or IL-1β, enhances STING-mediated immune responses by inhibiting the microtubule-mediated trafficking of STING from the Golgi apparatus to lysosomes for degradation ^30^. Additionally, a burst of mitochondrial reactive oxygen species (ROS), during inflammation, can trigger the opening of the mitochondrial permeability transition pore (mPTP), resulting in the extracellular release of mitochondrial DNA (mtDNA). Leakage of mitochondrial components into the cytosol acts as damage-associated molecular patterns (DAMPs), recruiting cGAS-STING and triggering an immune response ^31^. Indeed, the production of IL-1β induces the release of mtDNA, which activates cGAS-STING and enhances IFN-I production ^32^.

The role of the STING/IFN-I pathway in nociception has been a subject of debate over the past years. While administration of STING agonists through the central route causes acute analgesia in naive animals, peripheral STING agonists or IFN-I injection promotes mechanical allodynia ^11,14,33^. Furthermore, STING activation in response to exogenous ligands, acutely reduces mechanical allodynia in various neuropathic pain models through IFN-I/IFNAR1 signaling ^11,12^. To investigate the role of IFN-I specifically in neurons, we used a gain-of-function approach by expressing human N154S STING mutation in TRPV1 nociceptors, resulting in constitutive IFN-I production. Our findings indicate that IFN-I/IFNAR1 signaling regulates thermal but not mechanical sensitivity in both naive and inflammatory conditions. These results support previous studies highlighting the importance of TRPV1 neurons in thermal sensitivity and hyperalgesia. Accordingly, ablation of TRPV1 neurons or their central terminals in the spinal cord using chemical or genetic methods, reduces thermal sensitivity, while mechanical sensitivity remains unaffected, even after sensitization by inflammation or nerve injury ^34,35^. Additionally, chemogenetic inhibition of TRPV1 neurons using the Designer Receptors Exclusively Activated by Designer Drugs (DREADD) system does not alter mechanical responses ^36^. Notably, Yu and colleagues have identified two peptidergic populations of nociceptors in human DRG that co-express TRPV1 and TRPA1, but not PIEZO ^37^, suggesting that, at least in humans, TRPV1 neurons predominantly respond to noxious thermal and chemical stimuli, while their sensitivity to mechanical stimuli might be relatively lower.

Strikingly, while the development of thermal hypersensitivity was prevented in mice with nociceptor-specific hSTING mutation, the oedema caused by CFA was slightly enhanced. Gain-of-function mutations in STING, including the N154S mutation, underlie a type I interferonopathy called SAVI (STING-associated vasculopathy with onset in infancy). SAVI is a severe disease characterized by early-onset systemic inflammation, skin vasculopathy, and interstitial lung disease (ILD). SAVI patients exhibit high levels of C-reactive protein ^38^ and upregulated NF-kB related protein (e.g. IL-6) ^39^. Neuronal overproduction of IFN-I may directly impact the immune system in the periphery. IFN-I has been shown to indirectly inhibit tumor growth by acting on immune cells. Natural killer (NK) cells, the first line of defense against infections and tumors, depend on type I IFNs for maturation, activation, and homeostasis in the tumor microenvironment (TME) ^40,41^. IFN-I also promotes dendritic cell maturation and migration toward lymph nodes, where their activity is enhanced through the cross-presentation of tumor-associated antigens to CD8+ T cells ^42–44^. Additionally, IFN-I may enhance a pro-inflammatory environment by inhibiting Treg proliferation, function, and recruitment, as observed in the TME ^45–47^. Further studies are needed to elucidate the relative role of neuronal IFN-I in immune responses to inflammation.

Our findings also highlight IFNAR1 in the STING-IFN pathway. IFNAR is widely expressed in peptidergic nociceptors and their central terminals in the spinal dorsal horn ^14,48^. The localization of IFNAR1 in the presynaptic terminals suggests a role for STING-IFN-I in regulating neurotransmitter release. Indeed, Liu and colleagues reported that IFNAR activation inhibits excitatory synaptic transmission in somatostatin-positive excitatory neurons, by suppressing glutamate release from presynaptic terminals ^48^. Although IFNAR1 expression in nociceptors is prominent, non-neuronal IFNAR1 may also contribute to the anti-nociceptive effects of IFN-I. Emerging evidence suggests that spinal cord microglia and astrocytes play important roles in the regulation of pain hypersensitivity following tissue and nerve injuries ^49,50^. Intrathecal administration of poly I:C, a synthetic dsRNA, increases the expression of IRF-7 and IRF-9, indicating that both astrocytes and microglia respond to IFN-I ^51,52^. Nevertheless, using nociceptor-specific AAVs to knockdown IFNAR1 in TRPV1^cre^-GOF mice, we found that neuronal type I IFN suppressed nocifensive behaviors through their cognate IFNAR1 receptor on TRPV1 nociceptors.

Furthermore, comparative transcriptomic profiling of DRGs revealed a marked interferon signature associated with downregulated genes and upregulated ISGs in response to STING activation. Among the ISG repertoire, known anti-viral and anti-tumor genes were identified, including OASL, EIF2AK2 (PKR), Irf7, Ifit1, Ddx58, Usp18 and ISG15. Interestingly, the eukaryotic initiation factor 2 (eIF2), a key effector of cellular stress responses, participates in the reduction of protein translation via phosphorylation by PKR. This reduction in protein translation allows cells to conserve energy and modify gene expression to effectively manage stress conditions^53^. While exogenous IFN-I does not result in PKR-eIF2α activation in DRG ^14^, there is evidence suggesting the involvement of this pathway in the regulation of thermal sensitivity ^14,54^. Neurons may utilize the PKR-eIF2a pathway to modulate ion channel expression, including TRPV1 and Kv4 channels, and facilitate the resolution of inflammatory hyperalgesia.

One of the ISGs identified in TRPV1^cre^-GOF DRGs was the Kcnip1 gene, which encodes the KChIP1 auxiliary subunit of voltage-gated potassium (Kv) channels. In line with our findings, Kcnip1 is a candidate gene involved in heat pain and has been identified to be enriched in TRPV1+ neurons ^55–57^. KChIPs and transmembrane dipeptidyl peptidase-like proteins (DPPs) are part of the Kv4 channel complex that generates A-type K+ currents ^58–60^ and inhibits nociceptor excitability below the threshold for action potential generation^61^. KChIP1 modulates the trafficking, voltage-dependence, and inactivation kinetics of Kv4, thereby affecting the firing patterns and excitability of sensory neurons ^58,62,63^. While Kv4.1 channels are expressed in all DRG neurons, Kv4.3 is predominantly found in small-diameter neurons ^64–66^, specifically non peptidergic IB4+ and peptidergic TRPV1+ neurons but not CGRP+ neurons ^64,66–68^. Notably, Kv4.3 channels are distributed across the entire cell surface of sensory neurons, including their peripheral endings in the skin ^69,70^. Given that Kv4.3 operates within the subthreshold voltage range, this channel may function in the cutaneous endings to inhibit action potential initiation in response to non-noxious stimuli. Additionally, Kv4.3 exhibit high temperature sensitivity within a narrow hyperpolarized voltage range ^71^, a property that could potentially be influenced by the level of KChIP1 expression. Our findings demonstrate that the constitutive activation of STING and the production of IFN-I specifically reduce the excitability of TRPV1 neurons while leaving the properties of IB4+ neuron unchanged. This supports a model in which IFN-induced KChIP1 expression reduces action potential initiation in TRPV1+ nociceptors, thereby increasing the activation threshold of nociceptors to noxious stimuli.

Overall, our results describe a nociceptor-specific antiviral pain-resolving ISG response, which alters the expression of ion channel-associated protein KChIP1 and nociceptive ion channels such as TRPV1. This STING-mediated response helps counteract inflammatory hyperexcitability and restores the nociceptor threshold after inflammation. This modulation may occur through the PKR-eIF2a pathway, which has been previously described to control proteostasis in nociceptors^14,54^. While recent work has explored the role of SING in pain sensitivity within the context of neuropathic pain, our findings shows that inflammation leads to an upregulation of STING in nociceptors. This upregulation is associated with a type I IFN response that promotes resolution of inflammatory pain. Furthermore, our study provides important mechanistic insights into the ion channel and gene expression changes relevant to STING activation in the context of inflammation. Further studies are needed to elucidate the downstream signaling pathways and transcriptional changes that mediate the effects of IFN-I in nociceptors. Additionally, investigating the role of IFN-I/IFNAR1 signaling in other types of sensory neurons, such as non-peptidergic nociceptors, and in different pain models will be crucial for understanding the broader implications of this signaling pathway in pain resolution.

By identifying STING-ISGs as a key pain-resolving response that tunes down nociceptor excitability in inflammatory pain conditions, we have added an important innate immune molecule to the core list of pattern recognition receptors (PRRs) that regulate the integration of pain signals in primary afferent neurons ^12,72–75^. Harnessing the power of IFN-I and ISGs to alleviate hyperexcitability of nociceptors may provide a unique therapeutic arsenal for managing inflammatory pain conditions.

## Materials and Methods

### Mice

Age-matched 6-to 8-week-old littermate mice of both sex were used for experiments. C57BL/6 and B6.Rosa26-TdTomato reporter mice (ai14 strain) were purchased from Jackson Laboratory (Jax strain #007914). Nav1.8-cre transgenic (Tg) mice (also known as SNS-Cre mice) were from Rohini Kuner’s lab (U. Heidelberg, Germany) and previously published ^18^. Nav1.8-cre Tg mice were bred with ai14 mice to generate Nav1.8-cre/Tdtomato mice. All animal experiments were approved by the institutional animal care and usage committee (IACUC) at Boston Children’s Hospital and conducted according to institutional animal care and safety guidelines at Boston Children’s Hospital and Harvard Medical School. The STING-/- [Tmem173gt C57BL/6J] and the flox-GOF [B6N.CAG-loxP-STOP-loxP-constitutively-active-hSTING-N154S mutant-IRES-TdTomato] mice were provided by Dr. F. Jirik (University of Calgary), the TRPV1-cre and TRPV1-pHluorin mice ^20^ were bred at the University of Calgary Animal Resource Center. All mice were genotyped with the primers reported in **Table S2**. To study the gain of function of STING in TRPV1 neurons, TRPV1-Cre+/− mice were bred with flox-STING-hN154S mice to generate TRPV1 neurons expressing GOF (TRPV1-Cre+/-;hSTING-N154S+/- or TRPV1^cre^-GOF) mice and control (TRPV1-Cre-/-;hSTING-N154S+/- or GOF) littermates. All mice were housed under standard conditions with both drinking water and food available *ad libitum*. All experiments were conducted according to the protocols approved by the University of Calgary Animal Care Committee (AC19-0169) and the guidelines of the Canadian Council on Animal Care.

### Treatment with chemicals, antibodies and viral constructs

#### Complete Freund’s Adjuvant injections

Mice were anesthetized by isoflurane and injected in the right hind paw with 20 µl of Complete Freund’s Adjuvant (CFA, purchased from Sigma) using a Hamilton syringe fitted with a 30-gauge needle. Paw oedema was measured with an electronic caliper.

#### AAV injections

Adeno associated virus (AAV2-PHP.S) was used for expression of IFNAR1-shRNAs1-3 in TRPV1^cre-^GOF cKI animals. AAV2-CW3SL-CAG-DIO-IFNAR1-shRNA1-3-eGFP and AAV2-CW3SL-CAG-DIO-IFNAR1-scr-eGFP (2-5E13 GC/mL) were produced at the University of Calgary, Alberta, Canada. The shRNAs were designed using the SplashRNA algorithm ^76^. Newly born pups (postnatal day 5) were administered 10 µl (intra-peritoneal) of the viral construct at 1E13 GC/mL using a Hamilton syringe connected to a 30G syringe tip (Becton Dickinson). Pups were returned to their parent cages and weaned after 21 days.

#### Neutralizing antibodies

Anti-mouse IFN-β neutralizing antibody (300 ng/mouse, PBL Assay Science, 32400-1), rabbit polyclonal IgG control (300 ng/mouse, Biolegend, CTL-4112), anti-mouse monoclonal IFNAR1 neutralizing antibody (MAR1, I-401; 300 ng/mouse,Leinco Technologies, I-401) and mouse IgG control (300ng/mouse, Leinco Technologies, I-536,) were dissolved in sterile PBS and administered intrathecally. Briefly, mice were anesthetized by isoflurane, the dorsal fur of each mouse was shaved, the spinal column was arched, and a 30-gauge needle was inserted into the subarachnoid space between the L4 and L5 vertebrae. Intrathecal injections of 10 µl were delivered over a period of 5 seconds.

#### Blocking peptide

The TAT-KChIP1 blocking peptide (YGRKKRRQRRR-AWLPFARAAAIGWMPVA) was purchased from LifeTein. TAT-KChIP1 was dissolved in sterile saline and administered intrathecally (5 and 10 µg in 5ul). For control experiments, the TAT-KChIP1 peptide (10 ug) was denaturated at 100°C for 30 min.

## Behavioral tests

### Evaluation of thermal sensitivity (Hot plate test)

Thermal sensitivity was assessed by using the hot plate test (Bioseb, Pinellas Park, FL). Mice were placed on a metal hot plate set to 52°C ± 0.5°C. The latency from the moment the mouse was placed on the heated surface until it displayed the first overt behavioral sign of nociception, such as lifting or licking a hind paw, vocalization, or jumping, was measured. A timer was used to record the response time, and the mouse was immediately removed from the hot plate after responding or after a maximum cutoff of 30 seconds to prevent tissue damage.

### Evaluation of thermal hyperalgesia (Hargreaves test)

Thermal hyperalgesia was examined as previously described ^20^ by measuring the withdrawal latency of both the right (ipsilateral, i.l.) and left (contralateral, c.l.) hind paws using a focused beam of radiant heat (IR = 30) from a Plantar Test apparatus (UgoBasile). Briefly, mice were individually placed in a small, enclosed testing arena on top of a Plexiglas floor and allowed to acclimate for at least 90 minutes. The Plantar Test apparatus was positioned beneath the animal, so that the radiant heat was directed to the plantar surface of the hind paw. Three trials were performed for each mouse, with a cut-off time set at 15 seconds to prevent tissue damage.

### Evaluation of mechanical hyperalgesia (Von Frey)

Mechanical sensitivity was evaluated using the von Frey test by the method of Dixon ^77^ and adapted by Chaplan et al. ^78^. Mice were placed individually in small, enclosed testing arenas on top of a wire grid platform and were allowed to acclimate for a period of at least 90 min before paw withdrawal threshold measurement (PWT). Stimulation was applied using the up-and-down method. Calibrated von Frey filaments, in the range 0.02–1.4 g (Cat no. 58011, Stoelting, IL USA), were applied perpendicularly to the right hindpaw with sufficient force to cause a slight buckling against the paw for 3–5 s. A positive response corresponds to a paw withdrawal, flinching, or licking. The PWT was determined as previously described ^78^.

### Non-evoked pain behavior measurements

Non-evoked pain-related behaviors were assessed using the noninvasive animal behavior recognition system LABORAS (Metris, Hoofdorp, Holland). LABORAS is a fully automated behavior recognition and tracking system using mechanical vibrations to classify different natural behaviors (e.g., eating, drinking, climbing, locomotion, etc.) and has previously been validated for pharmacological studies ^79^. Mice were placed in the LABORAS cages for a period of 24 h, with drinking water and food available *ad libitum* and under light/dark cycles.

### Mouse DRG neurons

DRG neurons were harvested from mice and enzymatically dissociated in HBSS containing 2 mg/ml collagenase type I and 4 mg/ml dispase (both from Invitrogen) for 45 min at 37 °C. DRGs were rinsed twice in HBSS and once in Neurobasal A culture medium (Thermo Fisher Scientific) supplemented with 2% B-27, 10 % heat-inactivated fetal bovine serum (HI-FBS), 100 μg/ml streptomycin, 100 U/ml penicillin, 50 ng/ml of Nerve Growth Factor (NGF) and 50 ng/ml glial cell-derived neurotrophic factor (GDNF) (all from Invitrogen). Individual neurons were dispersed by trituration through a fire-polished glass Pasteur pipette in 4 ml media and cultured overnight at 37 °C with 5% CO2 in 95% humidity on glass coverslips previously treated with Poly-Ornithine and Laminin (both from Sigma-Aldrich).

### Electrophysiological recordings

DRG neurons were placed in a 1 ml volume chamber and continuously perfused at 1.5 ml/min with extracellular solution containing (in mM): NaCl 140, KCl 5, CaCl_2_ 2, MgCl_2_ 1, Hepes 10, D-glucose 10, for current clamp recordings. NaCl was replaced by choline for voltage clamp experiments. Recordings were conducted at room temperature on TdTomato+ neurons identified using 20x magnification on an inverted epi-fluorescence microscope (Olympus IX51, Olympus America Inc.). For the identification of the IB4 positive neurons, coverslips were treated with the plant lectin IB4 conjugated to fluorescein isothiocyanate (IB4-FITC, 10 µg/ml, Invitrogen, I21411) for 30 min. Whole-cell patch clamp experiments were performed in both voltage-clamp and current clamp mode using a software-controlled Axopatch 200B amplifier in combination with a Digidata 1550A digitizer. Data were low-pass filtered at 5 kHz before being sampled at 10 kHz using Clampex 11 (all from Molecular Devices). Borosilicate glass pipettes (Harvard Apparatus Ltd.) were pulled and polished to 2-3 MΩ resistance with a DMZ-Universal Puller (Zeitz-Instruments GmbH). Patch pipettes contained (in mM): K-gluconate 105, KCl 30, MgCl_2_ 4, EGTA 0.3, HEPES 10, Na_2_ATP 4, Na_3_GTP 0.3 and Na-phosphocreatine 10, pH 7.35. All chemicals were purchased from Sigma-Aldrich.

Under voltage clamp control, cells were held at −70 mV and membrane capacitance (Cm) and membrane resistance (Rm) were calculated in response to a -10 mV hyperpolarizing pulse using the Membrane Test function of Clampex. Resting membrane potential (Vm), rheobase (minimum current injection required to elicit an action potential) and resultant action potential amplitude of the neurons were evaluated by switching to current clamp mode, prior to returning to voltage clamp control. Access resitance (Ra) and holding current (Ih) were monitored throughout the experiments to ensure recording stability, and recordings with an unstable Ih or poor Ra (Initial Ra > 20 MΩ or Ra >10% change over the period of recording) were discarded. Data were analyzed from recordings with 75-95% compensation to minimize offsets due to large voltage-clamp errors.

### Data analysis, Statistical comparisons

Data were analyzed with Clampfit 11 (Molecular Devices), Easy Electrophysiology (Easy Electrophysiology Ltd) and Prism 9 (GraphPad).

The peak conductance (G) of the potassium current was calculated as follows:

> G = Ip/(Vm-EK)

Ip is the peak outward potassium current, Vm is command voltage and Ek is the estimated reversal potential. Activation and inactivation plots were fitted to the Boltzmann relation:

> *f(V)* = *Gmax*/(1 + exp ((*V*_1/2_ - V)/k))

where *G*_max_ is the maximal conductance, *V*_½_ is the voltage at which activation or inactivation is half-maximal, and *k* is the slope factor (a positive number for activation, a negative one for inactivation). These parameters were determined from a fit to each experiment, then averaged together to give the mean values (± SEM). To generate averaged, normalised activation and inactivation plots, individual experiments were normalised to *G*_max_ and averaged together. These were refitted to the Boltzmann equation; estimates of *V*_½_ and *k* obtained from these fits differed from the means mentioned above by < 1 mV.

### Type I Interferon measurement

Supernatants of primary DRG neuron cultures were collected 16 hours after stimulation with ADU-S100 (1 µg/ml; Chemietek, CT-ADUS100). To deplete TRPV1 neurons, the primary DRG cultures were treated with Resiniferatoxin (1 µM, RTX, Alomone, R-400) for 15 minutes before ADU-S100 treatment. The concentration of Type I IFN was measured using a Mouse IFN 2-Plex Discovery Assay® (Eve Technologies). Protein concentration was quantified using a Bradford assay (Bio-Rad Laboratories) for normalization.

### Microarray analysis of Nav1.8Tg-TdTomato Neurons

#### FACS purification of neurons

24 hours after CFA injection into the footpad, lumbar (L4-L6) DRGs ipsilateral to or contralateral to the side of CFA injection were dissected from mice, digested in 1 mg/mL Collagenase A/2.4 U/mL Dispase II (enzymes from Roche), dissolved in HEPES buffered saline (Sigma) for 70 minutes at 37°C. For each subsequent microarray sample, DRGs from n=3 mice each were pooled. Following digestion, cells were washed into HBSS containing 0.5% Bovine serum albumin (BSA, Sigma), filtered through a 70 µm strainer, resuspended in HBSS/0.5% BSA, and subjected to flow cytometry. Cells were run through a 100 µm nozzle at low pressure (20 p.s.i.) on a BD FACS Aria II machine (Becton Dickinson, USA). A neural density filter (2.0 setting) was used to allow visualization of large cells and TdTomato+ cells were gated on for isolation. For subsequent RNA extraction, fluorescent neurons were FACS purified directly into 1 mL Qiazol (Qiagen). FACS data was analyzed using FlowJo software (TreeStar, OR USA). Flow cytometry was performed in the IDDRC Stem Cell Core Facility at Boston Children’s Hospital.

#### RNA Processing, Microarray Hybridization and Bioinformatics Analysis

Total RNA was extracted by sequential Qiazol extraction and purification through the RNeasy micro kit with on column genomic DNA digestion according to the manufacturer’s instructions (Qiagen, CA, USA). RNA quality was determined by Agilent 2100 Bioanalyzer using the RNA Pico Chip (Agilent, CA, USA). Samples with RIN>7, were used for subsequent analysis. RNA was amplified into cDNA using the Ambion WT expression kit for Whole Transcript Expression Arrays, with Poly-A controls from the Affymetrix GeneChip Eukaryotic Poly-A RNA control kit (Affymetrix, CA, USA). The Affymetrix GeneChip WT Terminal labeling kit was used for fragmentation and biotin labeling. Affymetrix GeneChip Hybridization control kit and the Affymetrix GeneChip Hybridization, wash, stain kit was used to hybridize samples to Affymetrix Mouse Gene ST 1.0 GeneChips, fluidics performed on the Affymetrix GeneChip Fluidics Station 450, and scanned using Affymetrix GeneChip Scanner 7G (Affymetrix). Microarray work was conducted at the Boston Children’s Hospital IDDRC Molecular Genetics Core facility. Affymetrix CEL files were normalized using the Robust Multi-array Average (RMA) algorithm with quantile normalization, background correction, and median scaling. Differentially expressed transcripts were illustrated using volcano plots, generated by plotting fold-change differences against comparison p-values or –log (p-values).

### Microarray analysis TRPV1-pHluorin Neurons

#### Flow Cytometry

On day 3 of CFA, both ipsilateral and contralateral DRG sample (L4-L6) from TRPV1-ecGFP mice were collected and digested separately. After digestion, cells were filtered through a 90 mm mesh (Sarstedt) and washed in PBS 1% FBS. Cells were analyzed on a FACS Aria II (BD Bioscience). After FACS sorting, RNA was extracted separately from GFP positive and GFP negative cells using a RNeasy Mini kit (Qiagen). The quantity of RNA was determined using a Nanodrop 2000c spectrophotometer (Thermo-Fisher Scientific).

#### RNA Processing, Microarray Hybridization and Bioinformatics Analysis

Three biological replicates for the GFP positive and GFP negative samples (∼50 ng/sample) were submitted to the Centre for Applied Genomics (Toronto, Canada). Here the quality of the samples was assessed using the Agilent Bioanalyzer 2100 with the RNA Pico chip kit (Agilent Technologies). RNA integrity number values between 6.5 and 7 were achieved. The expression profiling was performed according to the manufacturer’s instructions with Affymetrix GeneChip Mouse Gene 2.0 ST Array (Affymetrix). Primary data analysis was carried out with the Affymetrix Expression Console 1.4.1.46 software including the Robust Multi-array Average module for normalization. Gene expression data were log-transformed. A change was considered significant when the FDR-corrected p-value/q-value thresholds met the criterion q < 0.01 at fold changes > 2 (expression increments or declines larger than two).

### RNA sequencing analysis of TRPV1^cre^-GOF Neurons

#### RNA Processing and sequencing

Neurons**-**Lumbar (L4-L6) DRGs were dissected from naïve GOF (n = 4) and TRPV1^cre^-GOF mice (n = 4). Total RNA from was purified through the RNeasy micro kit with on column genomic DNA digestion according to the manufacturer’s instructions (Qiagen, CA, USA). RNA quality was determined by Agilent Technologies 5200 Fragment Analyzer (Agilent, CA, USA). Samples with RIN>7, were used for subsequent analysis. cDNA libraries were generated were generated using the NEBNext® Ultra™ II Directional RNA Library Prep Kit for Illumina® (New England Biolabs) on an Illumina NextSeq500.

#### RNA sequencing analysis

The quality control of reads was performed with FastQC, and all samples passed quality control. Reads were then aligned with STAR (v2.4.0j) ^80^ on the Mus musculus genome GRCm39.107 from EBI databases. Alignment was then converted with SAMtools (v1.16.1) ^81^, and genes counts were computed with HTSeq counts (v0.9.1) ^82^.

##### Differential Gene Expression analysis

Count matrix from HTSeq-counts were analyzed with DESeq2 (v1.34.0) ^83^. In order to compare samples, raw data were normalized with rlogTransformation function. A change was considered significant when the adjusted p-value (Benjamini-Hochberg) thresholds met the criterion p < 0.05 at fold changes > 2 (expression increments or declines larger than two).

### Analysis of published single-cell RNAseq

To analyze STING (TMEM173) expression in peripheral sensory neurons, the «Adolescent, Level 6, Taxonomy Level 2 Pns neurons» dataset from the mouse brain atlas (http://mousebrain.org/)^21^ was extracted and processed using Seurat package (Seurat version 4.1.0) (59) in R (version 4.1.1). Loom files were converted into Seurat object using the LoadLoom function from SeuratDisk package. Results were then processed with the classical Seurat pipeline. The expression matrix for each population was computed with AverageExpression function, and the heatmap was generated using the DoHeatmap function of the mean expression matrix of the list of gene of interest.

### Immunohistochemistry

Animals were perfused with phosphate buffered saline (PBS) to wash out blood and then perfused with 4% paraformaldehyde (PFA) (Electron Microscopy Science, Cat. No. 15713). Lumbar (L4-L6) DRGs and spinal cord were extracted and dehydrated overnight in 30% sucrose and embedded in optimal cutting temperature (OCT) solution (VWR International, Cat. No. 95057-838). Embedded tissues were sliced 10-μm thick onto SuperFrost slides (VWR International). The following antibodies were used in this study: chicken polyclonal anti-GFP (1:500, Invitrogen, A10262), rabbit polyclonal anti-GFP (1:500, Invitrogen, A10262), rabbit polyclonal anti-STING (1:100, Cell Signaling, 13647S), rabbit polyclonal anti-TRPV1 (1:250, Alomone, ACC-030), guinea pig polyclonal anti-TRPV1 (1:250, Alomone, ACC-030-GP), rat polyclonal anti-Substance P (1:500, Millipore, MAB356), IB4-coupled Alexa 594 (1:1000, Invitrogen, I21412), IB4-coupled Alexa 488 (1:1000, Invitrogen, I21411), goat polyclonal anti-GFRα3 (1:500, R&D Systems,VFU021721), goat polyclonal anti-GFRα2 (1:500, R&D Systems, AF429), donkey anti-chicken Alexa Fluor 488 (1:1000, Invitrogen, SAB4600031), goat anti-rabbit Alexa Fluor 488 (1:2000, Invitrogen, A11008), goat anti-rabbit Alexa Fluor 555 (1:2000, Invitrogen, A21428), chicken anti-goat Alexa Fluor 647 (1:2000, Invitrogen, Cat. No. A21469;), anti-rat Alexa Fluor 647 (1:2000, Cederlane, 712-605-153).

All sections were mounted using Aqua PolyMount (Polysciences). Confocal images were acquired on a Zeiss LSM 510 Meta confocal microscope and AxioCam HRm camera and analyzed using a 20× objective. Sections were imaged and analyzed using ImageJ. A minimum of three mice per group were analyzed.

### In situ hybridization

#### Human DRG neurons

RNAscope in situ hybridization multiplex chromogenic assay was performed as instructed by Advanced Cell Diagnostics (ACD). Snap frozen human DRG tissues were cryo-sectioned at 18 µm, mounted on SuperFrost Plus slides (FisherScientific) and stored at −80°C. The next day, slides were removed from the −80°C freezer and immediately washed with PBS (pH 7.4; 5 min, twice), fixed with 4% PFA-PBS, and then dehydrated in 50% ethanol (5 min), 70% ethanol (5 min) and 100% ethanol (5 min) at room temperature. Slides were pre-treated with H2O2 10 min at RT and washed twice in distilled water. Then, slides were submerged in 1X boiling RNAscope target retrieval reagent for 5 minutes. After target retrieval agent treatment, slides were transferred in distilled water and then washed in 100% ethanol. The slides were air dried briefly and then boundaries were drawn around each section using a hydrophobic pen (ImmEdge PAP pen; Vector Labs). When hydrophobic boundaries have dried, protease III reagent was added to each section and slides were incubated for 20 min. at 40°C in a HybEZ oven (ACD). This last step was repeated once, and slides were washed with distilled water before RNAscope assay. The RNAscope assay was performed according to the manufacturer’s instructions using a HybEZ oven (ACD). The probes used were Hs-TMEM173 (ACD, # 433541), Hs-Scn10a (ACD, #406291-C2). At the end of the process, hematoxylin counterstain was performed (30% Gill hematoxylin n°1; Sigma) and slides were mounted with Vectamount mounting medium (Vector Labs). Slides were imaged at 40x with Hamamastu Nanozoomer 2 slide scanner.

#### Mouse DRG neurons

Co-detection of RNAscope in situ hybridization multiplex v2 fluorescent assay (ACD, #323110) combined with immunofluorescence was performed on mouse DRG neurons. Mice were euthanized with isoflurane and then perfused with PBS followed by 4% PFA as described above. DRGs were extracted and placed in 4% PFA overnight at 4°C followed by 30% sucrose overnight at 4°C and then embedded in OCT and stored at −80°C. OCT sections were cut into 10μm slices onto SuperFrost slides and stored at −80°C for up to two months. On the day of the assay, slides were washed In PBS (5 min, twice, RT), baked for 30 min at 60°C and then post-fixed in pre-chilled 4% PFA in PBS for 15 min at 4°C. The tissue was then dehydrated sequentially in 50% ethanol (5 min, RT), 70% ethanol (5 min, RT), and 100% ethanol (2 min, twice, RT). The tissue was then treated with H2O2 (10 min, RT) (ACD, #322335), washed twice in PBS with 0.1% Tween-20 (PBS-T) (Sigma, #P1379) (5 min, twice, RT), and then a barrier was created around the tissue using a hydrophobic pen (Vector Labs, #H-4000). The primary antibody, diluted in Co-detection Antibody Diluent (ACD, #323160) was applied to each section and stored overnight at 4°C. The following day, the tissue was washed in PBS-T (2 min, twice, RT), and then submerged in 10% Neutral Buffered Formalin (NBF) (30 min, RT) for post-primary fixation. The tissue was then washed in PBS-T (2 min, 4 times, RT), RNAscope Protease Plus (ACD, #322331) was applied to the tissue for 30 min at 40°C. The RNAscope assay was then performed according to ACD’s instructions. At the end of the assay, before mounting the slides, the secondary antibody was added to the tissue for 30 min at RT before the slide was washed in PBS-T (2 min, twice, RT). The slides were then mounted using ProLong Gold antifade reagent with DAPI (Invitrogen, #P36931). The probes used were Mm-Trpv1 (ACD, #313331-C2), Mm-Kcnip1 (ACD, #466891-C3), Hs-TMEM173 (ACD, #433541), and Mm-Ifnar1 (ACD, #512971) and were labelled using TSA Vivid Fluorophores 570 (ACD, 323272) and 650 (ACD, #323273). The primary antibody used was anti-GFP (Chromotek, #PABG1) and the secondary antibody was Alexa Fluor 488 donkey anti-rabbit IgG (Invitrogen, #A21206). The tissue was imaged using the Leica SP8 confocal microscope at 20x or 63x objectives.

### Western Blot Analysis

#### TBK1 expression

To look at the phosphorylation of TBK1, we performed primary DRG neurons culture from WT and STING-/- mice and stimulated them with 10 µg/ml or 30 µg/ml ADU-S100 for one, three or six hours. DRGs were homogenized using a bullet blender (Next Advance) with SSB02 beads (Next Advance) and lysed in RIPA buffer with protease and phosphatase inhibitors (Thermo Scientific) for 45 minutes. Lysates were centrifuged at 10,000g for 10 min at 4°C, supernatants were collected, and protein concentration was quantified and normalized using a Bradford assay (Bio-Rad laboratories). Total lysates were separated by SDS-PAGE and transferred onto nitrocellulose membranes (Sigma-Aldrich). Membranes were blocked in 5% BSA 1h at room temperature, and then probed with anti-phospo-TBK1 antibody (1:200 dilution in 1% BSA, Cell Signalling, 5483S) at 4°C overnight. Membranes were then washed three times with TBS-T and incubated with horseradish peroxidase (HRP)-conjugated anti-rabbit antibodies (1:1000 in 1% BSA; Cederlane, NA934) for 1h at room temperature. Bands were visualized using the Immobilon Western chemiluminescent HRP Substrate (Bio-Rad Laboratories), and band density was calculated using Image J. Intensity of anti-TBK1 antibody (1:1000 dilution in 1% BSA; Cell signalling, 3504S) band was used for normalization among samples.

#### KChIP1 expression

DRGs were homogenized using a bullet blender (Next Advance) with SSB02 beads (Next Advance) and lysed in RIPA buffer with protease and phosphatase inhibitors (Thermo Scientific) for 45 minutes. Lysates were centrifuged at 10,000g for 10 min at 4°C, supernatants were collected, and protein concentration was quantified and normalized using a Bradford assay (Bio-Rad laboratories). Total lysates were separated by SDS-PAGE and transferred onto nitrocellulose membranes (Sigma-Aldrich). To investigate the level of KChIP1 protein, membranes were blocked in 5% BSA 1h at room temperature, and then probed with anti-KChIP1 antibody (1:250 dilution in 1% BSA, Alomone, APC-141.) at 4°C overnight. Membranes were then washed three times with TBS-T and incubated with horseradish peroxidase (HRP)-conjugated anti-rabbit antibodies (1:1000 in 1% BSA, Cederlane, NA934) for 1h at room temperature. Bands were visualized using the Immobilon Western chemiluminescent HRP Substrate (Bio-Rad Laboratories), and band density was calculated using Image J. Intensity of Rabbit anti-Beta-tubulin III antibody (1:1000 dilution in 1% BSA; Sigma-Aldrich, T2200) band was used for normalization among samples.

### Cytokine profiling

Blood samples were centrifuged (5000 x g, 30 minutes, 4°C) and then serum was processed using both a Mouse IFN 2-Plex Discovery Assay® and a MILLIPLEX Mouse Cytokine Array Proinflammatory Focused 10-plex (Eve Technologies).

### qPCR

DRGs were harvested and dissociated using a bullet blender (Next Advance) with SSB02 beads (Next Advance) in RLT buffer (Qiagen). Total RNA was extracted using a RNeasy Mini kit (Qiagen), according to the manufacturer’s instructions. The quality and quantity of RNA were determined using a Nanodrop 2000c spectrophotometer (Thermo-Fisher Scientific). Relative gene expression (normalized to GAPDH) was determined by qPCR using BlasTaq 2X qPCR MasterMix (ABMgood, G892) and a StepOnePlus real-time PCR detection system (Applied Biosystems). The designed primers for DNA amplification are listed in Supplemental **Table S3**.

## Data availability

Microarray data from lumbar DRG Nav1.8-cre Tg/TdTomato neurons from CFA ipsilateral and contralateral sides are deposited at the NCBI GEO database under accession number GSE221834. Microarray data collected from naïve lumbar DRG Nav1.8-Cre Tg/TdTomato neurons were previously deposited at GEO database under accession no. GSE46546. Microarray data collected from lumbar DRG TRPV1-ecGFP neurons from CFA ipsilateral and contralateral sides were previously deposited at GEO database under accession no. GSE201227. Bulk RNAseq data collected from lumbar DRG neurons from GOF and TRPV1^cre^-GOF animals were deposited at NCBI GEO database under accession no. GSE236865.

### Statistics

Statistical analyses were performed with GraphPad Prism 7^®^ software. Normal distribution was verified using D’Agostino-Pearson normality Test. For Gaussian data, Student’s t-test was used to assess statistical significance when comparing two means, One-Way ANOVA followed by the Tukey *post hoc* test was used to compare more than two groups and Two-way ANOVA followed by Bonferroni (two groups) and Tukey *post hoc* test (for more than two groups) for multiple comparisons. For non-Gaussian data, the non-parametric Mann Whitney U test was used to assess statistical significance when comparing two means, Kruskal-Wallis followed by the Dunn’s *post hoc* test was used to compare more than two groups. Statistical significance was established at P ≤ 0.05. Values were expressed as means ± standard error mean (SEM).

## Acknowledgments

This work was supported by operating grants from the Crohn’s and Colitis Canada (CCC) grant (CA) and The Canadian Institutes of Health Research (CIHR) grant # 388441 (CA). MD holds a Postdoctoral Fellowship Support from the Alberta Children’s Hospital Research Institute (ACHRI). FA was sponsored by the Eyes High fellowship initiative at U of Calgary. NA holds the Association of Gastroenterology (CAG) scholarship.

We would like to thank Dr. F. Jirik and Carolina Salazar for providing the STING-/- [Tmem173gt C57BL/6J] and the flox-GOF [B6N.CAG-loxP-STOP-loxP-constitutively-active-hSTING-N154S mutant-IRES-TdTomato] mice; and Frank Visser for producing the AAVs. We thank Dr. Clifford Woolf for generous support of this project and feedback. We thank the Animal care facility of U of Calgary for their assistance in mice care, the Clara Christie Centre for Mouse Genomics (CCCMG), the Flow Cytometry facility of the Snyder Institute and the Centre for Health Genomics and Informatics (CHGI).

---

**Supp Figure 1.**
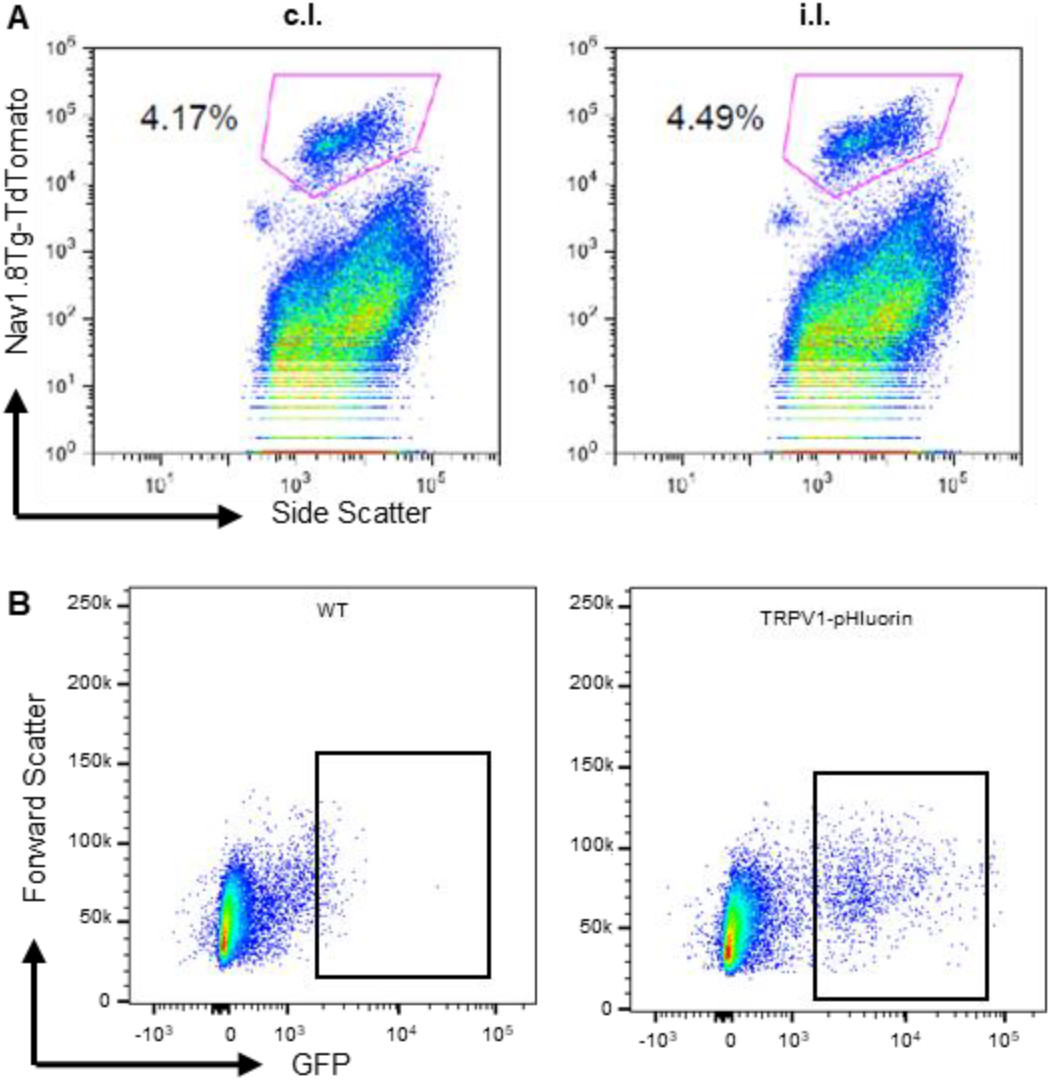
FACS gating strategy to isolate fluorescently labeled neurons from Nav1.8-cre Tg/TdTomato and TRPV1-ecGFP mice. **(A)** FACS plots are shown as representative example of the gating strategy used for Nav1.8-cre Tg/TdTomato lumbar DRG neuron isolation from contralateral and ipsilateral sides following CFA injection. Lumbar DRG were pooled from three mice per sample. (**B**) FACS plots are shown as representative example of the gating strategy used for WT and TRPV1-pHluorin DRG neurons isolation from ipsilateral sides following CFA injection. Lumbar DRG were pooled from three mice per sample.

**Supp Figure 2.**
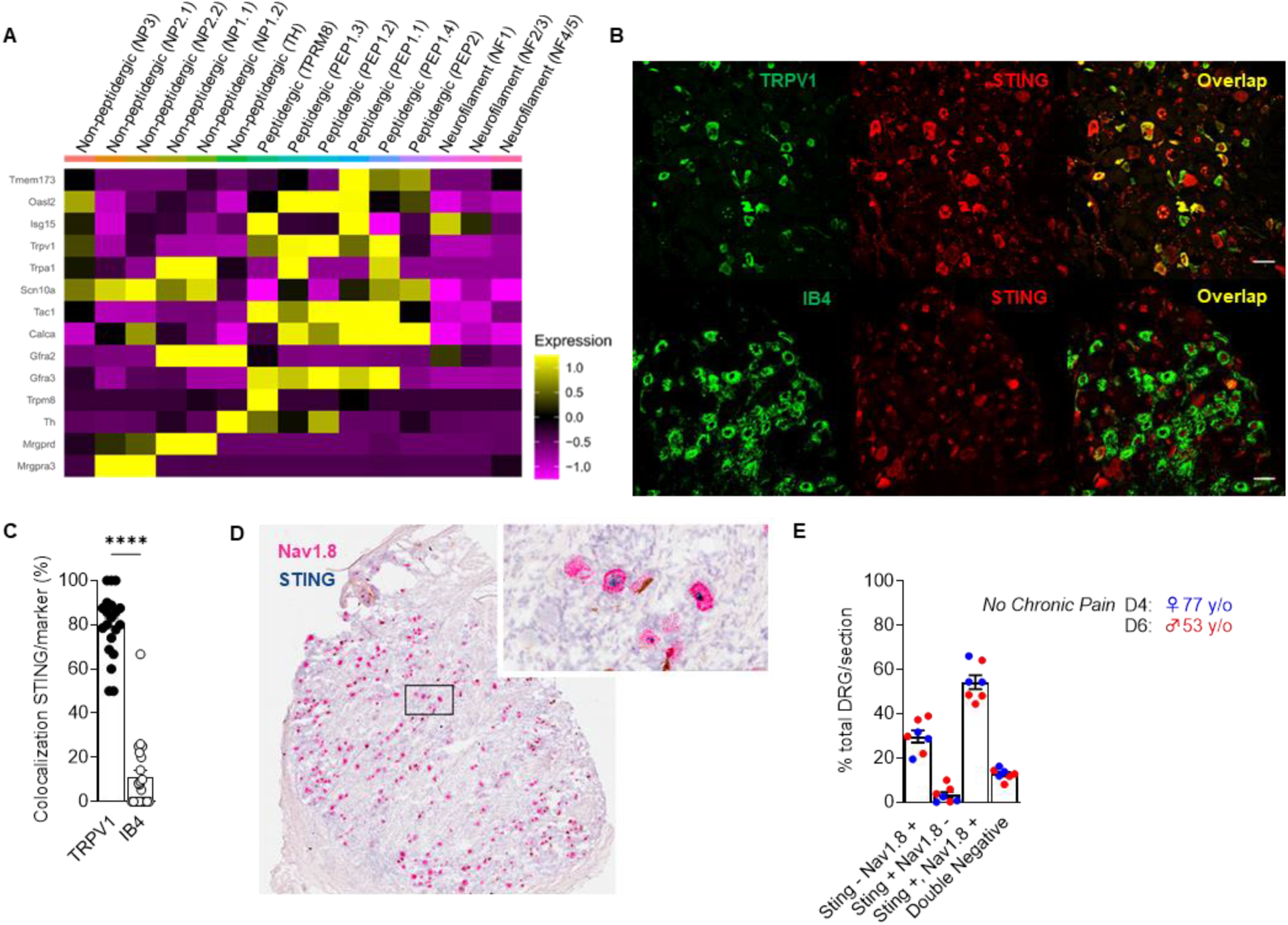
STING is expressed in mouse and human DRG nociceptors. **(A)** Heatmap of the expression of STING and selected population markers on the 15 populations of sensory neurons from DRG described in Zeisel et al., ^21^. **(B**) Representative confocal images of co-immunostaining for STING, IB4 and TRPV1 in mouse DRG neurons. Scale bars: 50 µm. (**C**) Dot plot showing the percentage of peptidergic (TRPV1) and non-peptidergic (IB4) neurons co-expressing STING in DRG from TRPV1-pHluorin mice (n = 3 mice). Statistical analysis was performed using T-test (**** p<0.001). (**D**) Representative RNAscope image showing expression of STING (light blue) in human DRG neurons co-expressing Nav1.8 (pink). (**E**) The bar graph summarizes the results (each symbol represents a single DRG; n = 3 for one patient and n = 4 for another patient). Results indicate the mean ± SEM.

**Supp Figure 3.**
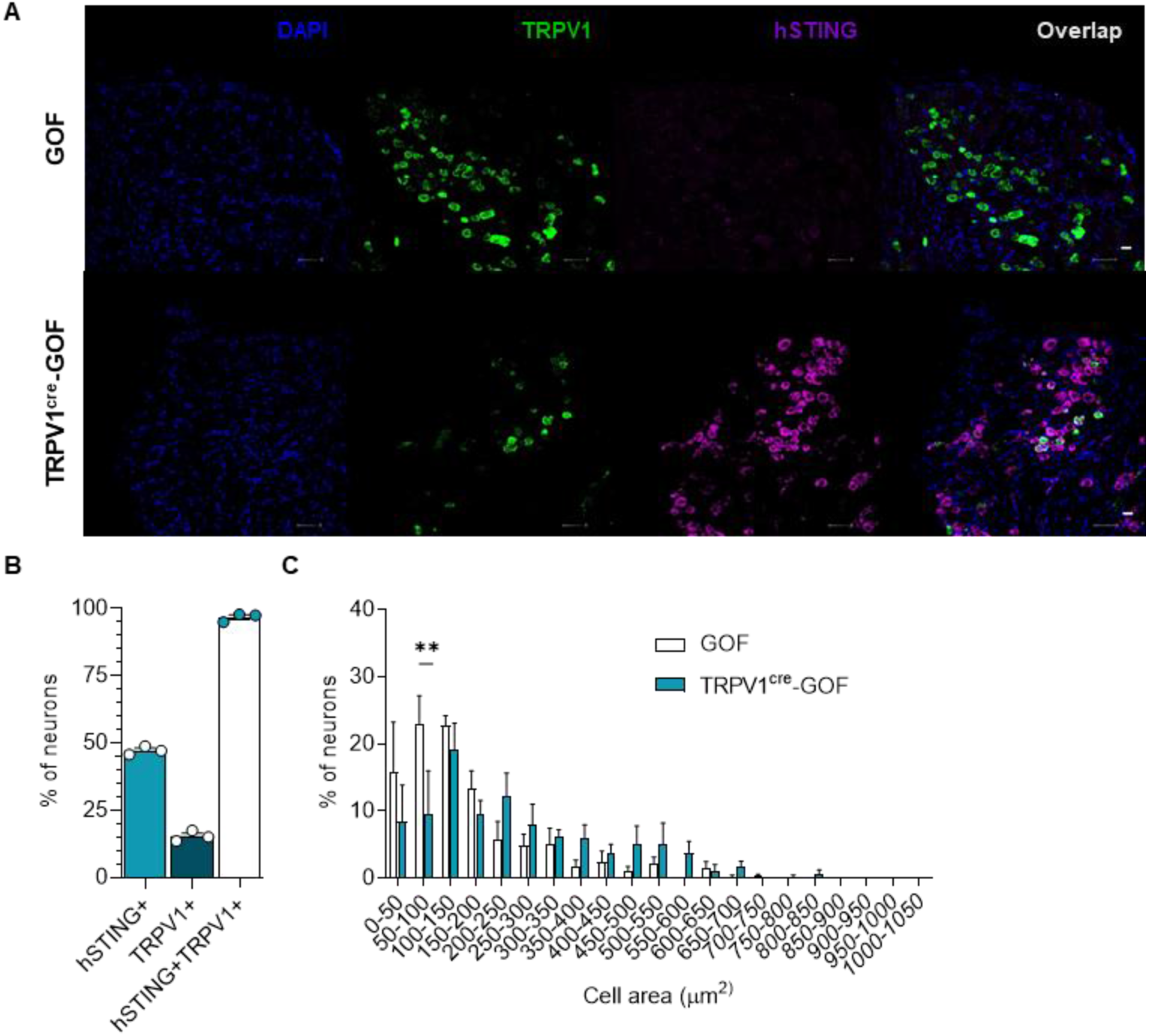
Nociceptor-specific hSTING-N154S mutation alters the numbers of TRPV1 neurons. (**A**) Expression of hSTING and TRPV1 in DRGs from TRPV1^Cre^-GOF and GOF littermate controls. Scale bars: 50 µm. (**B**) The bar graph summarizes the percentage of hSTING+ and TRPV1+ neurons normalized to the total number of neurons. Note that ∼100% of hSTING+ neurons are TRPV1+ neurons (each symbol represents a single animal n = 3). (**C**) Size distribution of the percentage of DRG neurons immunopositive for TRPV1, in naïve GOF (n = 3) and TRPV1^cre^-GOF mice (n = 3). Statistical analysis was performed using Two-way ANOVA followed by Tukey post-hoc test (** p < 0.01). Results indicate the mean ± SEM.

**Supp Figure 4.**
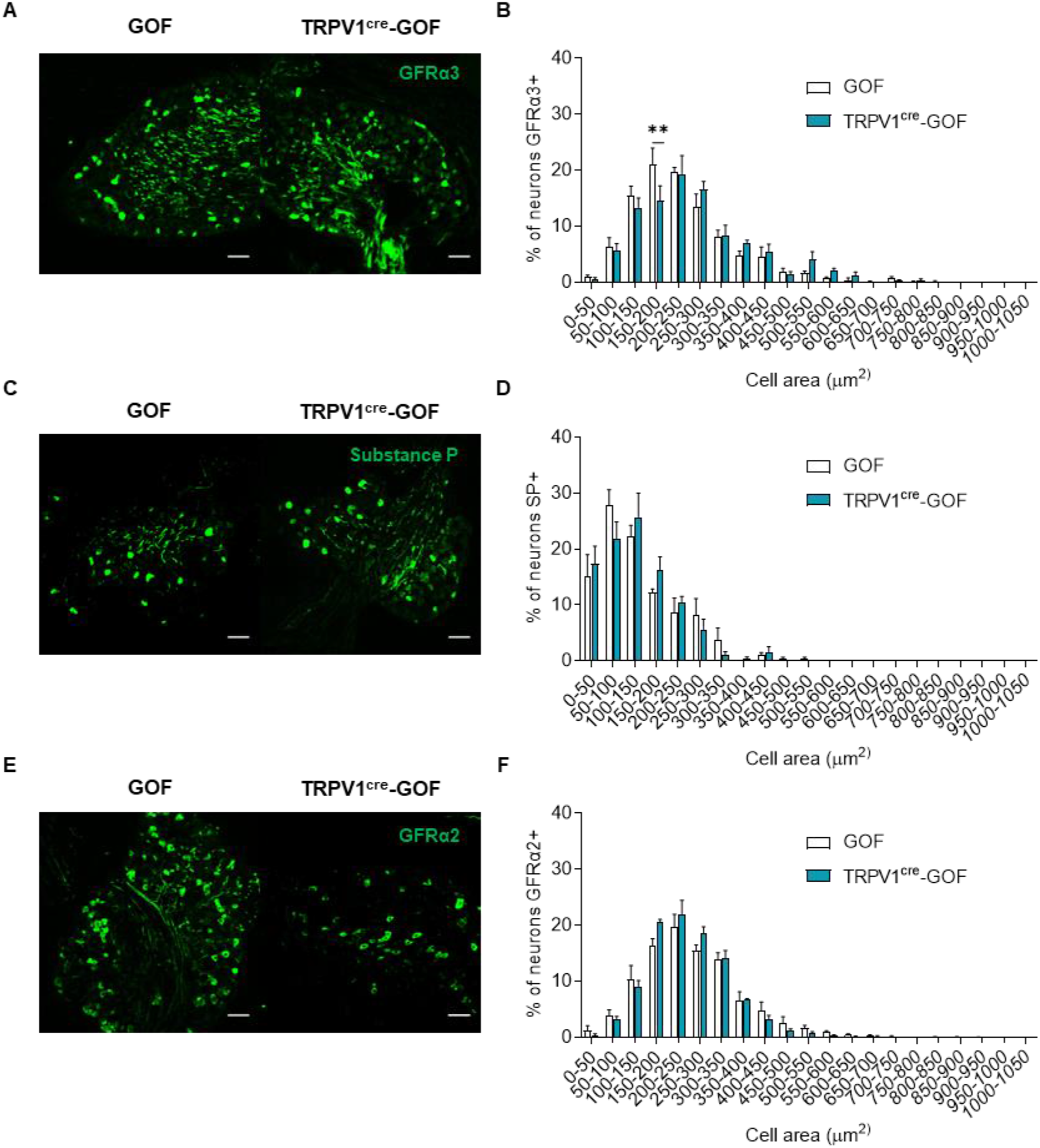
Characterization of peptidergic and non-peptidergic neurons in mice expressing the nociceptor-specific hSTING-N154S gain of function mutation. Representative confocal images of immunostaining for GFRα3 (A), Substance P (C) and GFRα2 (E) in DRGs of TRPV1^Cre^-GOF and GOF mice. Scale bars: 50 µm. Size distribution of DRG neurons positive for GFRα3 (B), Substance P (D) and GFRα2 (F) in TRPV1^Cre^-GOF (n = 3) and GOF mice (n = 4). Statistical analysis was performed using Two-way ANOVA followed by Tukey post-hoc test (** p < 0.01). Results indicate the mean ± SEM.

**Supp Figure 5.**
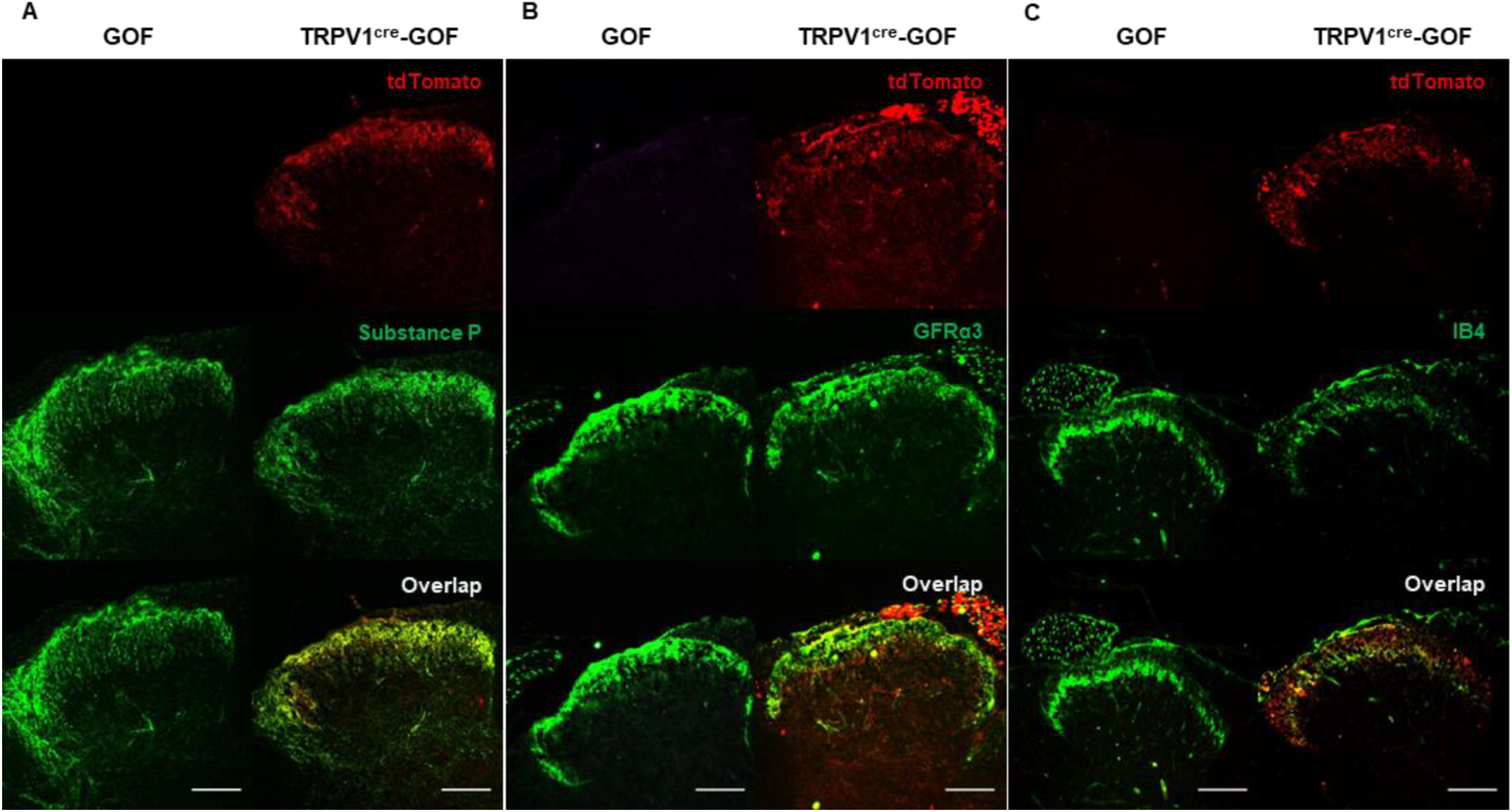
Nociceptor-specific hSTING-N154S mutation does not alter the neuronanatomical organization of TRPV1 neurons. Representative confocal images of lumbar spinal cord section from TRPV1^Cre^-GOF and GOF littermate controls, immunostained for Substance P (**A**), GFRα3 (**B**) and IB4 (**C**). Scale bars: 100 µm.

**Supp Figure 6.**
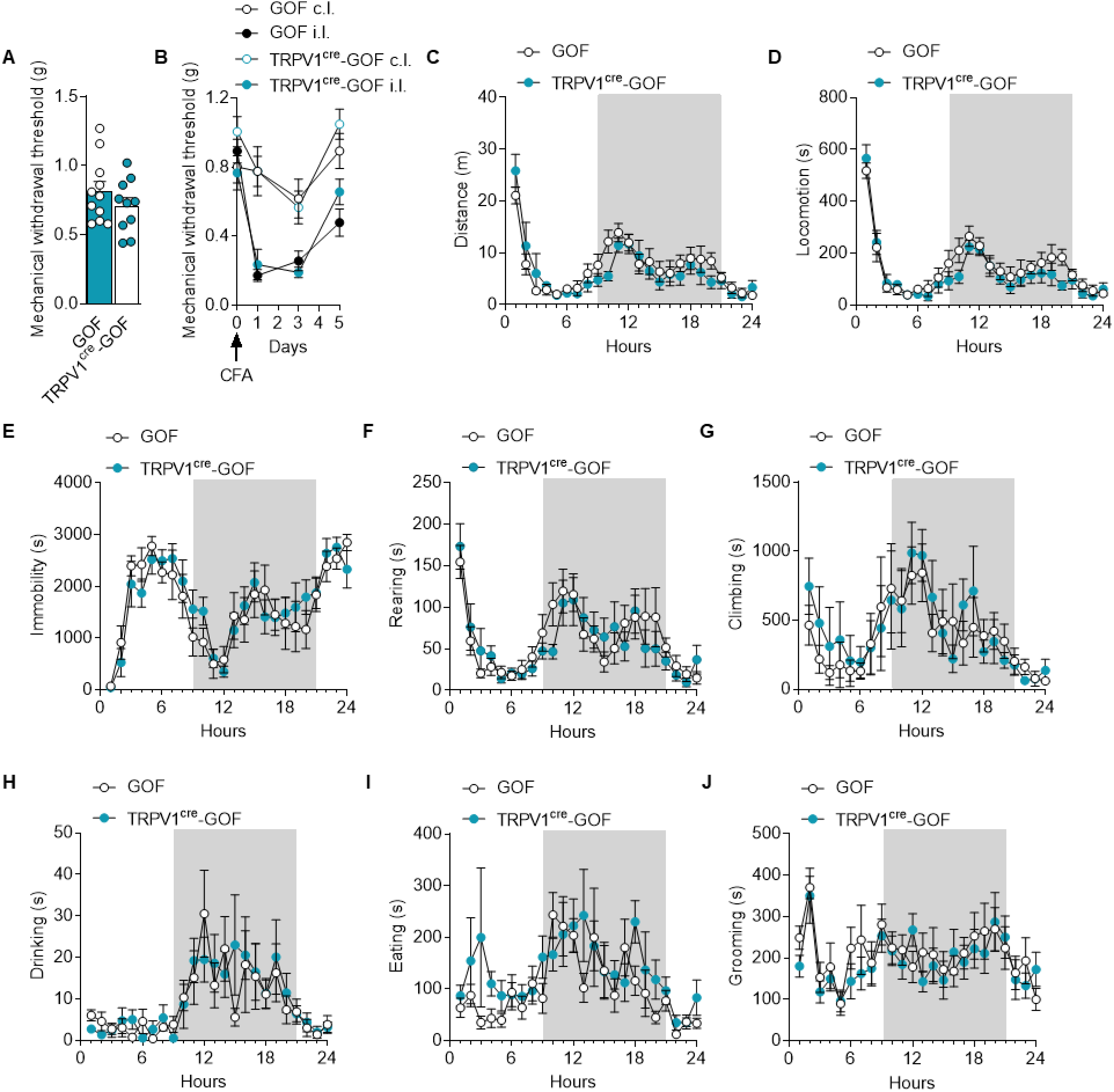
Mice expressing the nociceptor-specific hSTING-N154S mutation exhibit no changes in non-evoked pain-related behaviors and mechanical hyperalgesia. (A) Mechanical sensitivity in naïve TRPV1^Cre^-GOF and GOF littermate controls (n = 10 and 11, respectively). (B) Measurement of mechanical allodynia in TRPV1^Cre^-GOF and GOF littermate controls following intraplantar CFA (n = 6 and 8, respectively). (C-J) Non-evoked normal behaviors of both naïve GOF (n = 7) and TRPV1^cre^-GOF (n = 9) mice. Distance (C), Locomotion (D), Immobility (E), Rearing (F), Climbing (G), Drinking (H), Eating (I), Grooming (J) behaviors were measured using the Laboras system for 24 hours. Statistical analysis was performed using Two-way ANOVA followed by Tukey post-hoc test. Results indicate the mean ± SEM.

**Supp Figure 7.**
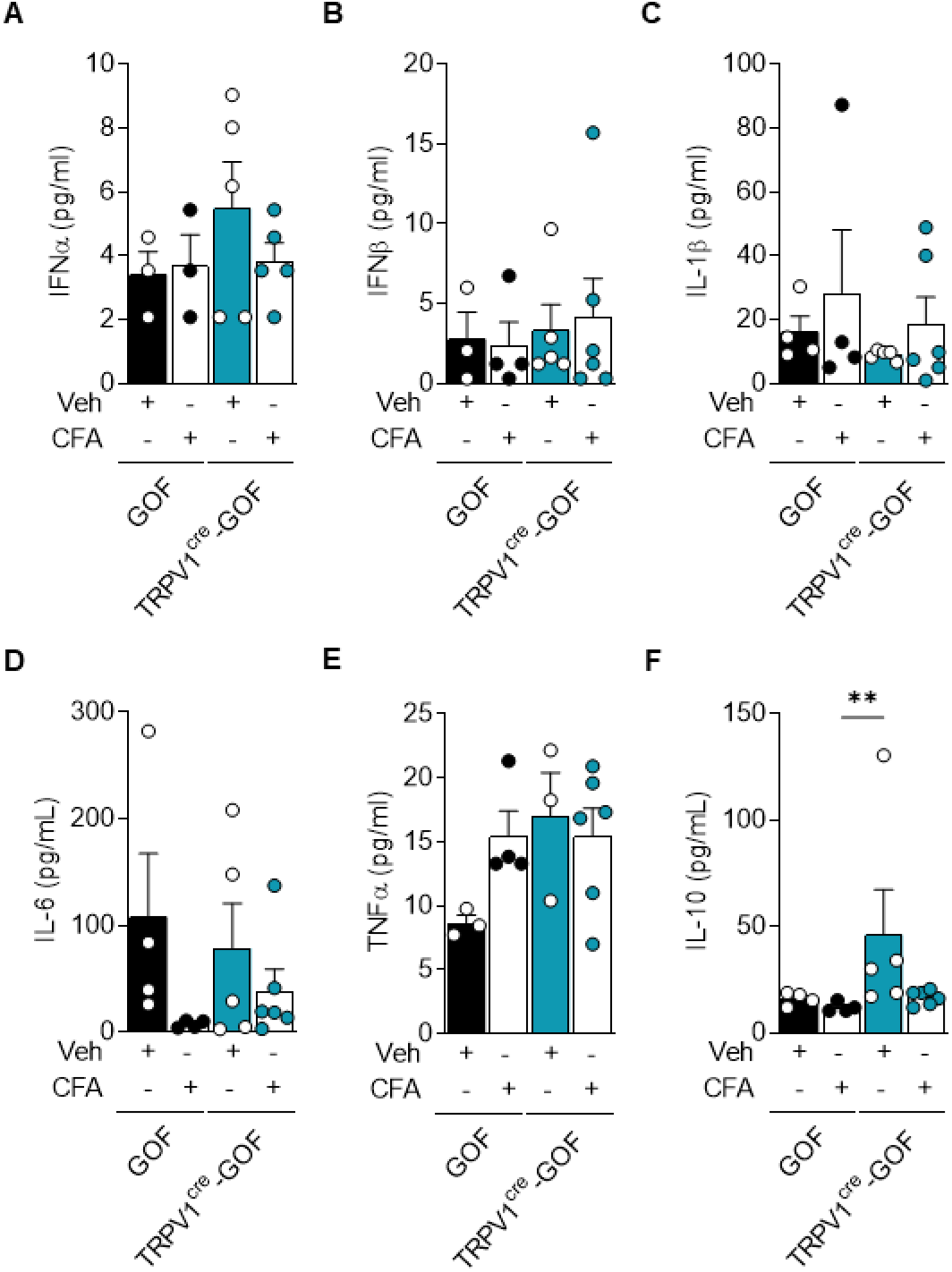
Nociceptor-specific hSTING-N154S mutation does not induce systemic inflammation. IFNα (**A**), IFNβ (**B**), IL-1β (**C**), IL-6 (**D**), TNF-α (**E**) and IL-10 (**F**) levels were determined by Luminex technology in the serum of GOF (n = 4), TRPV1^cre^-GOF (n = 5), CFA-treated GOF (n = 4) and CFA-treated TRPV1^cre^-GOF (n = 6) mice. Statistical analysis was performed using Kruskal-Wallis followed by Dunn’s post hoc test. Results indicate the mean ± SEM.

**Supp Figure 8.**
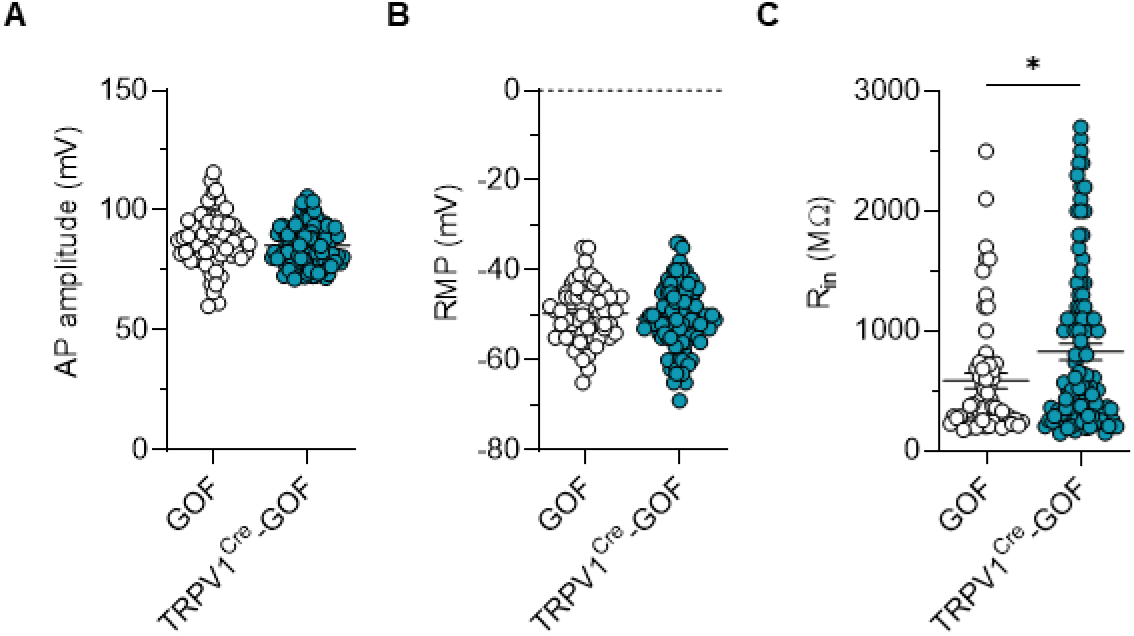
Electrophysiological properties of TRPV1-hSTING-N154S expressing neurons. **(A)** Measurement of action potential amplitude in TRPV1 neurons from TRPV1^Cre^-GOF mice and GOF littermate controls. **(B)** Measurement of the resting membrane potential of TRPV1 neurons from TRPV1^Cre^-GOF mice and GOF littermate controls. **(C)** Measurement of the input resistance of TRPV1 neurons from TRPV1^Cre^-GOF mice and GOF littermate controls. Results indicate the mean ± SEM (n=61 for GOF and n=101 for TRPV1^Cre^-GOF). Statistical analysis was performed using unpaired t-test (* p < 0.05). Results indicate the mean ± SEM.

**Supp Figure 9.**
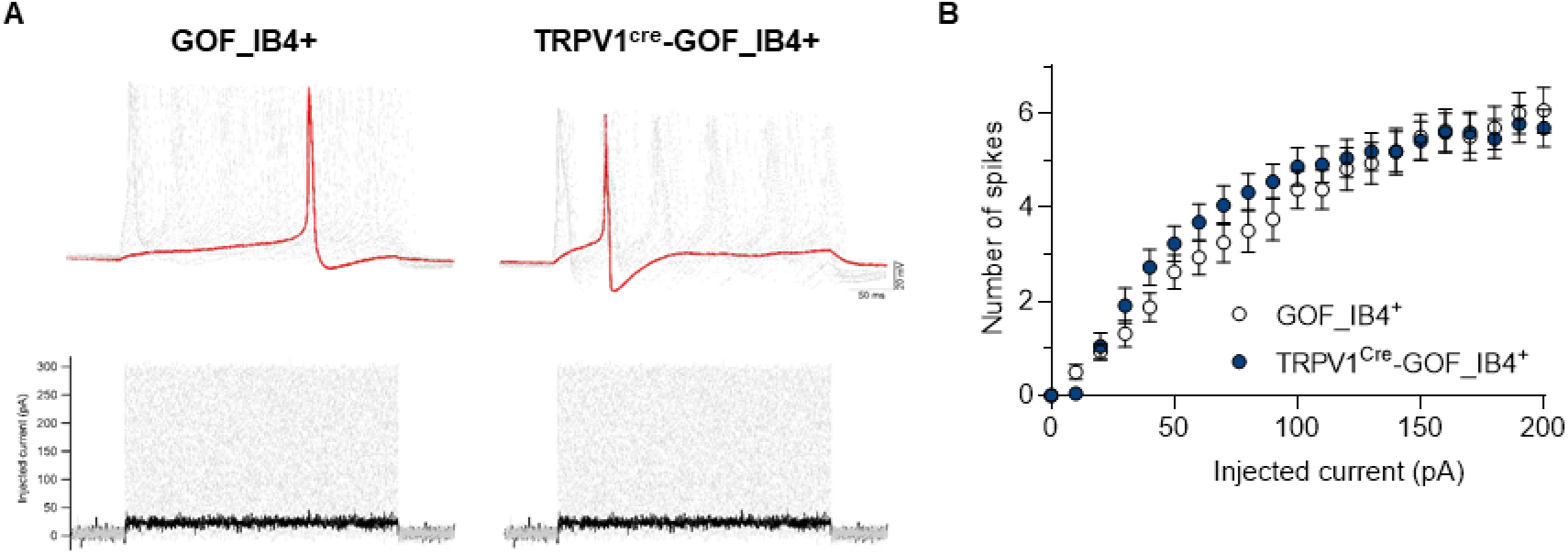
Nociceptor-specific STING-N154S gain of function mutation does not alter the excitability of non-peptidergic neurons. **(A)** Representative current clamp recording of IB4+ neurons from TRPV1^Cre^-GOF mice and GOF littermate controls. Cells were injected with 500 ms current pulses with an increment of 10 pA and an interval of 5 seconds. Scale bars: 20 mV/50 ms. **(B)** Number of spikes as a function of injected current in IB4+ neurons from TRPV1^cre^-GOF mice and GOF littermate controls. Statistical analysis was performed using a two-way ANOVA test with Tukey’s post-hoc test. Results indicate the mean ± SEM.

**Supp Figure 10.**
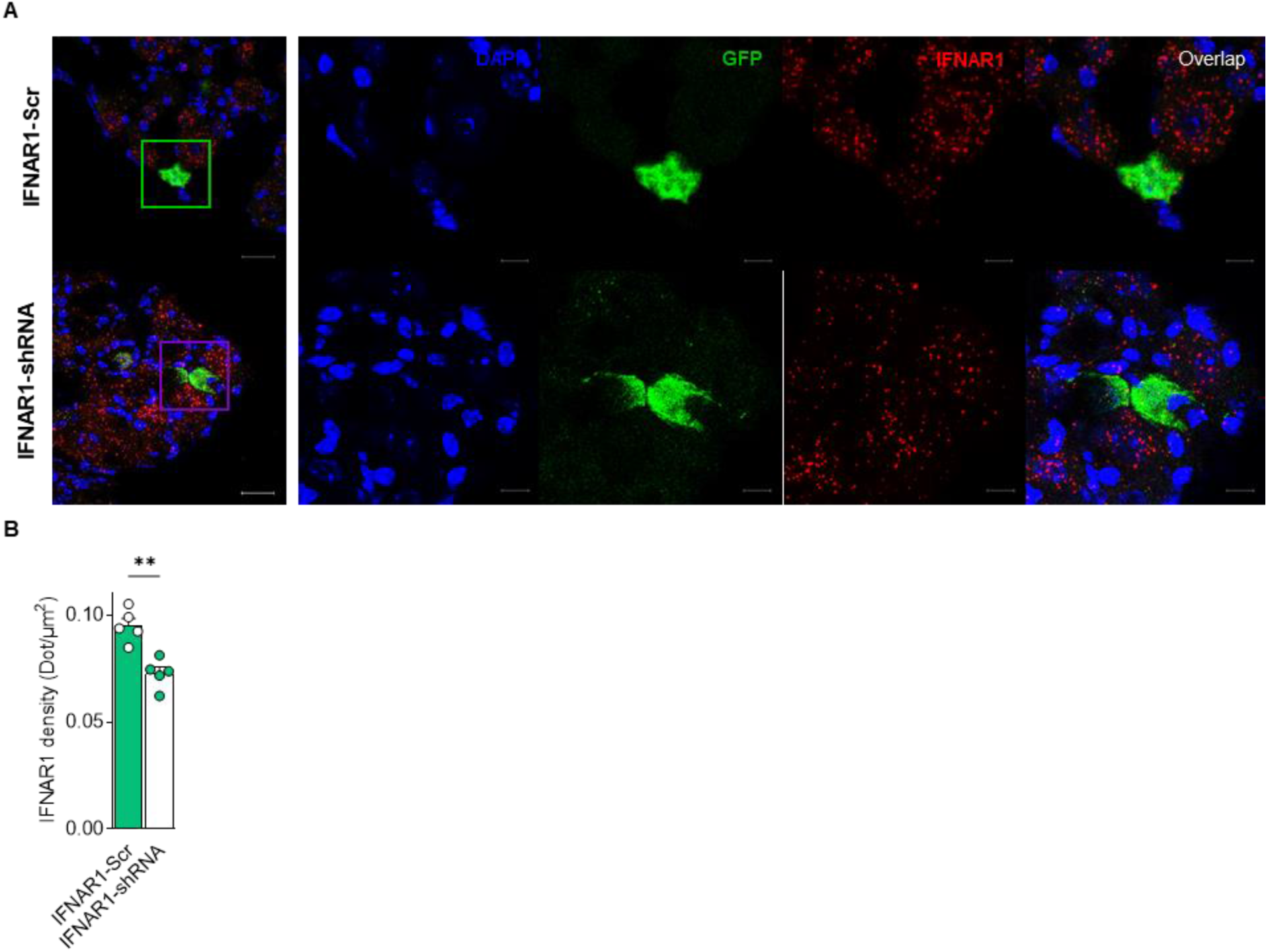
Knockdown of IFNAR1 in TRPV1 neurons from TRPV1^Cre^-GOF mice. **(A)** Representative confocal image of TRPV1 neurons from TRPV1^Cre^-GOF mice injected with AAV-DIO-Scr-shRNA (n = 5) or AAV-DIO-IFNAR1-shRNA (n = 5). Images represent DAPI staining, AAV-GFP expression (green) and IFNAR1 transcripts (red) by RNAscope. Scale bars: 25 µm and 10 µm on the cropped images. **(B)** Quantification of IFNAR1 density measured by the number of transcripts represented by dots per surface unit in AAV-infected TRPV1 neurons. Statistical analysis was performed using unpaired t-test (** p < 0.01). Results indicate the mean ± SEM.

**Table S1.** Cytokine profiling was determined by Luminex technology in the serum of GOF (n = 4), TRPV1^cre^-GOF (n = 5), CFA-treated GOF (n = 4) and CFA-treated TRPV1^cre^-GOF (n = 6) mice. Statistical analysis was performed using Kruskal-Wallis followed by Dunn’s post hoc test. Results indicate the mean ± SEM.

| Cytokine | GOF Control (pg/ml ± SEM) | GOF CFA (pg/ml ± SEM) | TRPV1<sup>cre</sup>-GOF Control (pg/ml ± SEM) | TRPV1<sup>cre</sup>-GOF CFA (pg/ml ± SEM) | P value |
| --- | --- | --- | --- | --- | --- |
| GM-CSF | 34.70 ± 3.228 | 26.16 ± 1.229 | 32.96 ± 2.764 | 28.81 ± 2.745 | 0.1114 |
| IL-2 | 20.78 ± 4.683 | 11.17 ± 1.127 | 10.19 ± 1.093 | 13.04 ± 1.072 | 0.0596 |
| IL-4 | 4.415 ± 1.244 | 1.500 ± 0.4823 | 3.448 ± 1.815 | 1.457 ± 0.2974 | 0.1088 |
| IL-12 | 95.55 ± 20.62 | 39.28 ± 6.904 | 239.6 ± 172.8 | 43.86 ± 6.527 | 0.1292 |
| MCP-1 | 334.3 ± 190 | 136.3 ± 15.44 | 322.2 ± 115.6 | 121.1 ± 11.25 | 0.2296 |
| IFN-γ | N.D. | N.D. | N.D. | N.D. | N.D. |

**Table S2.**
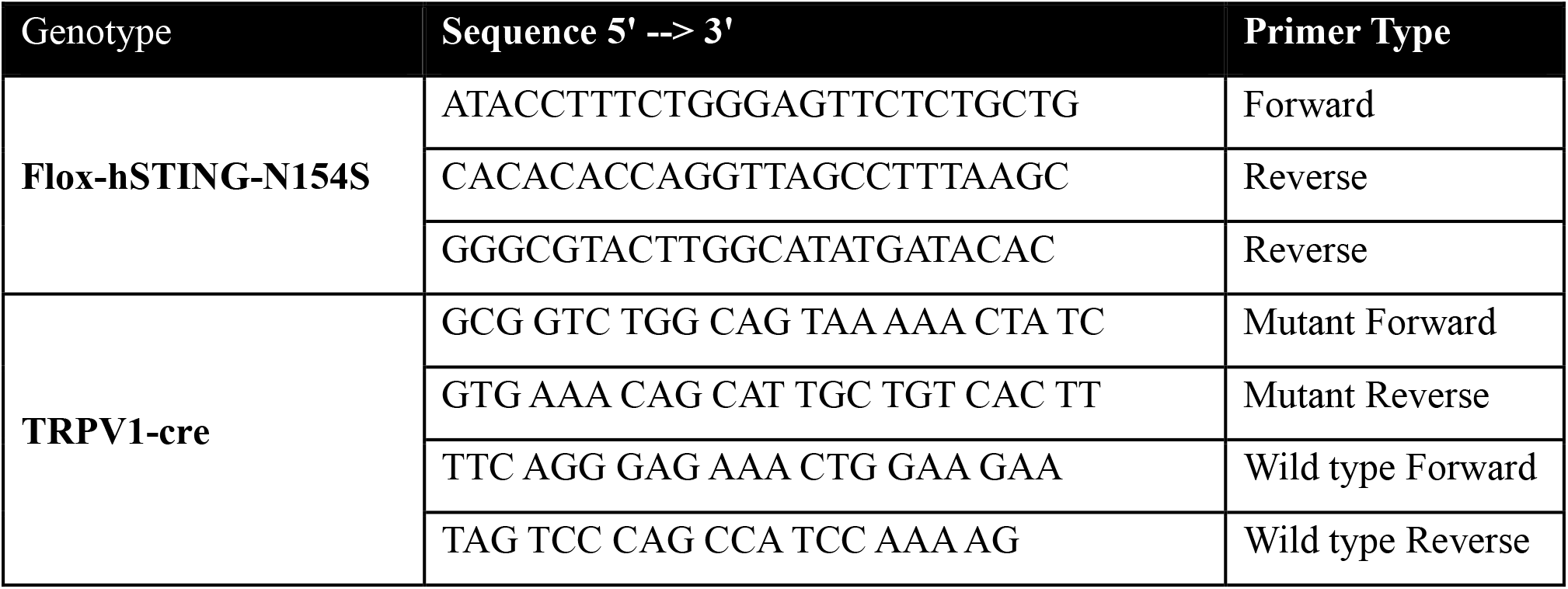
Primer sequences.

**Table S3.**
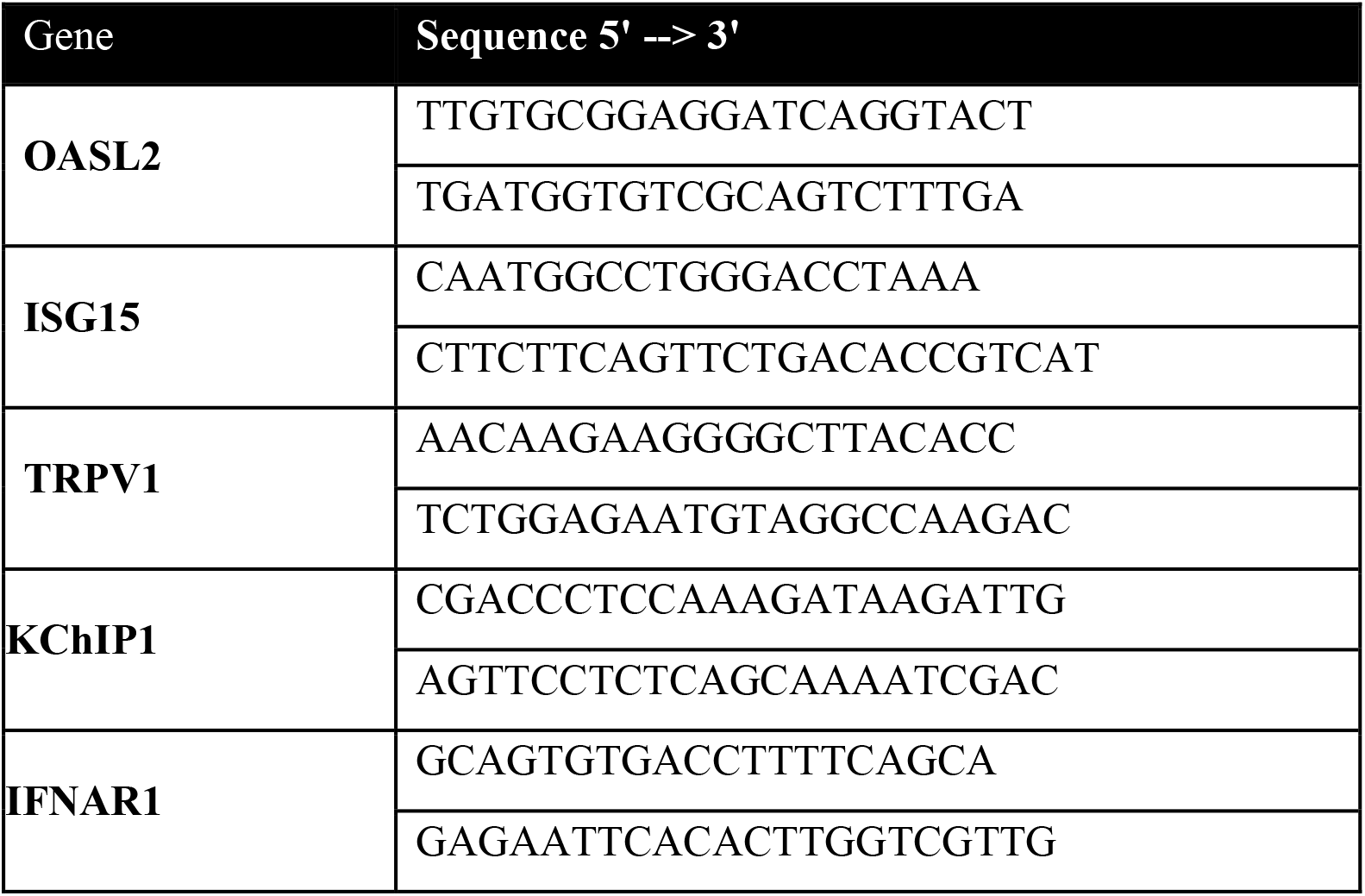
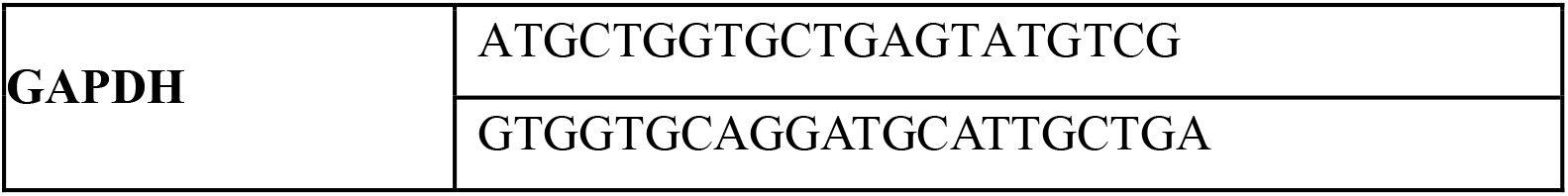
qPCR primer sequences.

